# Suppressing APOE4-induced mortality and cellular damage by targeting VHL

**DOI:** 10.1101/2024.02.28.582664

**Authors:** Wei I. Jiang, Yiming Cao, Yue Xue, Yichun Ji, Benjamin Y. Winer, Mengqi Zhang, Neel S. Singhal, Jonathan T. Pierce, Song Chen, Dengke K. Ma

**Author notes:** Correspondence (S.C.) & (D.K.M.).

## Abstract

Mortality rate increases with age and can accelerate upon extrinsic or intrinsic damage to individuals. Identifying factors and mechanisms that curb population mortality rate has wide-ranging implications. Here, we show that targeting the VHL-1 (Von Hippel– Lindau) protein suppresses *C. elegans* mortality caused by distinct factors, including elevated reactive oxygen species, temperature, and *APOE4*, the genetic variant that confers high risks of neurodegeneration in Alzheimer’s diseases and all-cause mortality in humans. These mortality factors are of different physical-chemical nature, yet result in similar cellular dysfunction and damage that are suppressed by deleting VHL-1. Stabilized HIF-1 (hypoxia inducible factor), a transcription factor normally targeted for degradation by VHL-1, recapitulates the protective effects of deleting VHL-1. HIF-1 orchestrates a genetic program that defends against mitochondrial abnormalities, excess oxidative stress, cellular proteostasis dysregulation, and endo-lysosomal rupture, key events that lead to mortality. Genetic *Vhl* inhibition also alleviates cerebral vascular injury and synaptic lesions in *APOE4* mice, supporting an evolutionarily conserved mechanism. Collectively, we identify the VHL-HIF axis as a potent modifier of APOE4 and mortality and propose that targeting VHL-HIF in non-proliferative animal tissues may suppress tissue injuries and mortality by broadly curbing cellular damage.

## INTRODUCTION

Age-related mortality is a universal phenomenon observed across all biological species. Understanding the factors that modulate this trajectory is essential for developing strategies to mitigate the impact of aging on population health. Intrinsic genetic determinants and host physiology, extrinsic environmental challenges and abiotic stress, as well as stochastic events all interact to confer mortality risks. In humans, genetic association studies have identified major genetic risk factors for all-cause mortality, including the ε4 allele of the *APOE* gene (*APOE4*)^1–4^. This allele also represents the highest genetic risk factor for late-onset Alzheimer’s disease (AD) as well as the highest genetic risk modifier of early-onset forms of AD^5–7^. Emerging human studies implicate *APOE4* homozygosity as a major genetic cause, not just a risk modifier, of AD that constitutes one of the most frequent human Mendelian disorders^8^. APOE4 proteins differ in cholesterol transport capabilities compared to its allelic counterparts and, contrary to its heightened association with AD risk, it is linked to decreased susceptibility to age-related macular degeneration^9–11^. Genetic variations including non-*APOE4* variant alleles of *APOE* have also been shown to be associated with reduced mortality in rare long-lived human centenarians^12^. These studies have provided intriguing cases of how genetic variations may link to mortality and age-related diseases in humans. However, despite these advances, establishing causal and mechanistic relationships among genetic variations, cellular processes, environmental impacts, and mortality rates at the population level remains a formidable challenge.

To identify causal genetic factors that influence mortality and to elucidate their underlying mechanisms, the nematode *Caenorhabditis elegans* emerges as a well-suited model organism. Its amenability to genetic manipulation, short lifespan, and well-characterized genome provide an ideal platform for discovering novel genetic modifiers of age-related mortality and pathologies within the context of a whole organism and with well-controlled environmental conditions^13–15^. In addition, the relatively simple and transparent anatomy of *C. elegans* allows for direct observation of cellular and physiological changes throughout its lifecycle, facilitating the identification of cellular mechanisms and their impact on mortality and pathologies. Pioneering investigations of longevity mutants in *C. elegans* have underscored the importance of the insulin, PI3K and mTOR pathways, leading to discoveries of their evolutionarily conserved roles governing the aging process across various eukaryotic organisms, including humans^16–18^. Besides the trajectory of aging under normal culture conditions, *C. elegans* is also subject to rapidly increased mortality when exposed to severe environmental stresses, including elevated temperature, pathogen infection and abiotic stress^15,19–21^. While mild stress can extend longevity through the mechanism of hormesis^22,23^, it remains largely unknown how mortality accelerates when *C. elegans* is severely stressed.

Genetic studies in *C. elegans* have identified reduction- or loss-of-function (LOF) alleles, including those of *daf-2* and *vhl-1*, which can extend longevity and confer broad stress resilience^24–27^. *daf-2* encodes a homolog of insulin receptors that orchestrate anabolic metabolism, autophagy regulation, and somatic maintenance program during aging. *daf-2* mutants are exceptionally long lived and stress resistant. *vhl-1*, the ortholog of the Von Hippel–Lindau tumor suppressor gene, encodes an E3-ubiquitin ligase that targets the hypoxia-inducible factor HIF-1 for degradation. Loss of VHL-1 stabilizes HIF-1 and activates a genetic program linked to both longevity extension and stress resilience. While HIF-activating VHL mutations in humans increase risks to various cancers, including clear cell renal cell carcinoma, HIF and its target gene activation in non-proliferative cells, such as neurons and cardiomyocytes, can be protective against ischemic insults, reperfusion injuries and metabolic stress^28–30^. Although previous transcriptomic and proteomic studies unveiled many transcriptional targets of HIF, specific mechanisms underlying the protective effect of the VHL-HIF axis in the context of mortality, longevity regulation and stress resilience still remain unclear.

In light of the escalating mortality rates associated with aging and exacerbated by diverse intrinsic and extrinsic factors, our study aimed to identify factors and mechanisms capable of mitigating these outcomes. We find that *vhl-1* loss or stabilized HIF-1 strongly suppresses *C. elegans* population mortality induced by diverse factors, including elevated reactive oxygen species (ROS), temperature stress, and transgenic expression of the human AD-neurodegenerative and longevity risk variant *APOE4*. We identify functional HIF-1-regulated genes that may contribute to guarding against cellular processes mechanistically linked to age-related mortality. We further used *APOE4*-humanized mice to highlight the likely evolutionarily conserved mechanism by which VHL inhibition mitigates the APOE4 effects on animal tissue injury and mortality.

## RESULTS

### Roles of VHL-1 in suppressing mortality

We showed previously that transgenic neuronal expression of human *APOE4*, but not *APOE3,* in *C. elegans* exacerbated neurodegeneration^31^. To study potential effects of APOE4 on population mortality using a fast, reproducible and robust model, we examined the mortality trajectory (lifespan curve) of *APOE4*-transgenic *C. elegans* under various constant conditions of temperature stress beyond the normal range (15 °C to 25 °C). When subjected to a constant temperature of 28 °C, wild-type animals died within a few days (median lifespan of approximate 4 days post L4), whereas neuronal *APOE4* expression drastically shortened the lifespan (median lifespan fewer than 2 days post L4) (Fig. 1a). Under such constant heat stress, *APOE4* expression also led to profound morphological deterioration of the PVD neuron (Fig. 1b).

**Figure 1.**
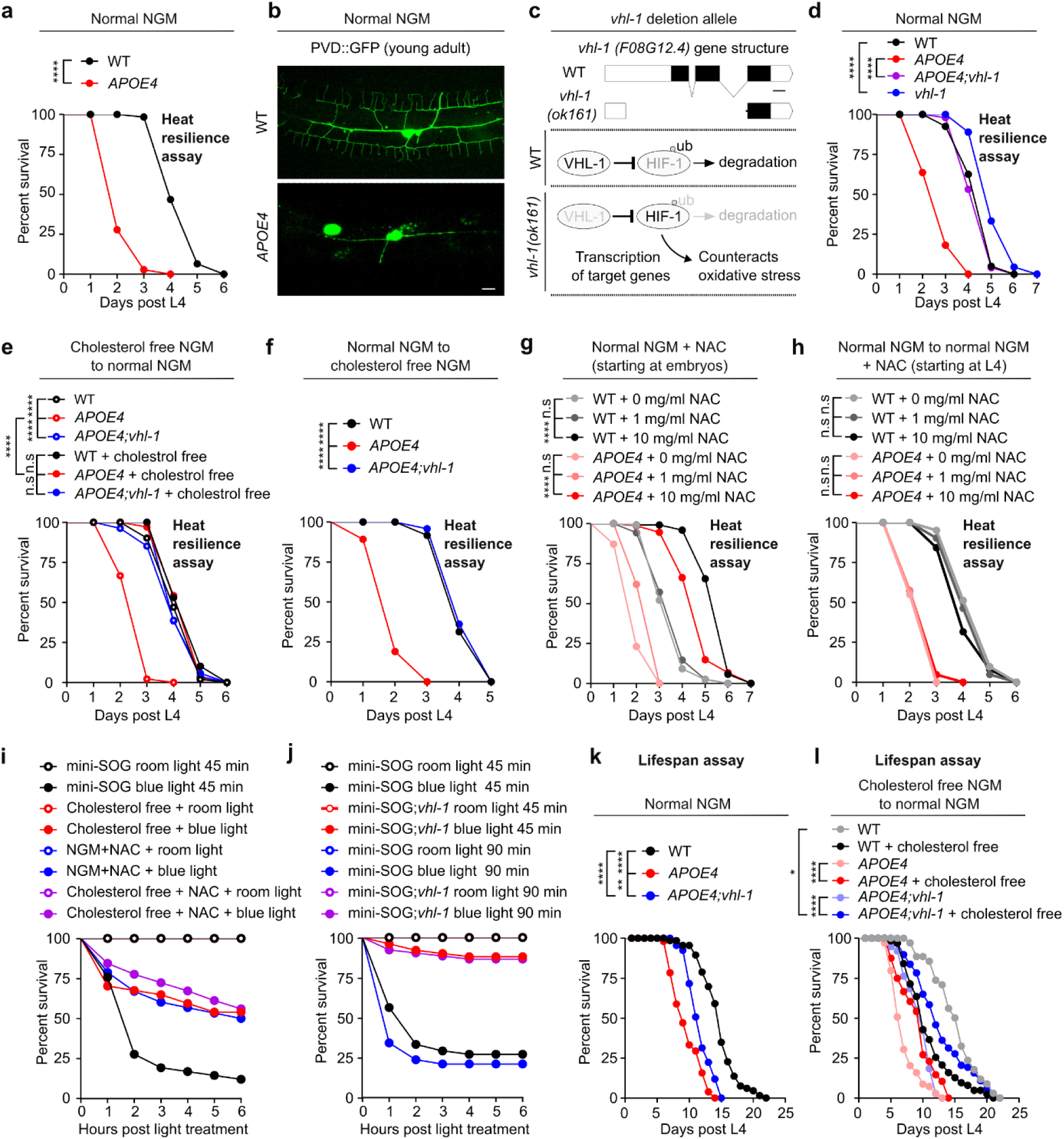
Loss of *vhl-1* suppresses mortality induced by multiple factors (miniSOG, heat and *APOE4*) that cause molecular and cellular damages. (a) Lifespan curves of N2 wild type (WT) and pan neuronal *APOE4(vxIs824)* transgenic animals at 28 ℃ starting at L4 on normal NGM, showing 50% median and 50% maximal survival decrease in *APOE4(vxIs824)* compared to WT. **** indicates P < 0.0001 (WT: n=62 animals, *APOE4*: n= 36 animals). (b) Representative confocal microscopic images of PVD neuron (*wyIs592[ser-2prom-3p::myr-GFP]*) in WT and pan neuronal *APOE4(vxIs824)* animals at young adult stages on normal NGM, showing PVD abnormalities with apparent loss of third and fourth branches. Scale bar: 10 μm. (c) Schematic of *vhl-1(ok161)* loss-of-function deletion allele (with the exon 2 and 3 deleted) that leads to impaired ubiquitination and stabilized HIF-1 to counteract oxidative stress. Scale bar: 100 bp. (d) Lifespan curves of WT, pan neuronal *APOE4(vxIs824)*, *vhl-1(ok161*) mutants, and *APOE4(vxIs824)*; *vhl-1(ok161*) animals at 28 ℃ starting at L4 on normal NGM. **** indicates P < 0.0001 (WT: n=40 animals, *APOE4*: n= 44 animals, *APOE4(vxIs824)*; *vhl-1(ok161*): n=49 animals, *vhl-1*: n= 45 animals). (e) Lifespan curves of WT, *APOE4 (vxIs824)* and *APOE4(vxIs824)*; *vhl-1(ok161*) with or without early life (starting at embryos) cholesterol-free NGM to L4 on cholesterol-free NGM followed by picking to normal NGM and culturing at 28℃. **** indicates P < 0.0001, n.s indicates non-significant (WT: n=51 animals, *APOE4*: n= 45 animals, *APOE4(vxIs824)*; *vhl-1(ok161*): n=54 animals, WT + cholesterol free: n=49 animals, *APOE4* + cholesterol free: n=35 animals, *APOE4(vxIs824)*; *vhl-1(ok161*) + cholesterol free: n=52 animals). (f) Lifespan curves of WT, *APOE4*(*vxIs824*) and *APOE4(vxIs824)*; *vhl-1(ok161*) grown to L4 on normal NGM followed by picking to cholesterol-free NGM and culturing at 28℃. **** indicates P < 0.001 (WT: n=48 animals, *APOE4*: n= 37 animals, *APOE4(vxIs824)*; *vhl-1(ok161*): n= 47 animals). (g) Lifespan curves of WT and *APOE4(vxIs824)* starting at early life (starting at embryos) with indicated NAC diet concentration (0 mg/ml, 1 mg/ml and 10 mg/ml) to L4 on normal NGM supplemented with indicated NAC concentration followed by picking to normal NGM supplement with indicated concentration of NAC and culturing at 28℃. **** indicates P < 0.001, n.s indicates non-significant (WT+ 0 mg/ml: n=340 animals, WT+ 1 mg/ml: n=357 animals, WT+ 10 mg/ml: n=122 animals, *APOE4* + 0 mg/ml: n=78 animals, *APOE4* + 1 mg/ml: n=29 animals, *APOE4* + 10 mg/ml: n=74 animals). (h) Lifespan curves of WT and *APOE4(vxIs824)* grown to L4 on normal NGM followed by picking to normal NGM supplemented with indicated concentration of NAC (starting at L4) and transferred to 28℃. *** indicates P < 0.001, n.s indicates non-significant (n > 40 animals per condition). (i) Percent survival of miniSOG animals [*unc-25p::tomm20::miniSOG::SL2::RFP*], grown to L4 starting at early life (embryos) with NAC supplement, starting at early life (embryos) with cholesterol free NGM or normal NGM, followed by room light or blue light treatments for 45 mins. (n > 40 animals per condition). (j) Percent survival of miniSOG animals [*unc-25p::tomm20::miniSOG::SL2::RFP*] or LOF mutant *vhl-1(ok161)*; miniSOG animals grown to L4 on normal NGM, followed by room light or blue light treatments for 45 mins or 90 mins (n > 40 animals per condition). (k) Lifespan curves of WT, *APOE4(vxIs824)*, *APOE4(vxIs824)*; *vhl-1(ok161*) animals at constant 20 ℃ on normal NGM. ** indicates P < 0.01, **** indicates P < 0.0001, (n > 40 animals per condition). (l) Lifespan curves of WT, *APOE4(vxIs824)*, and *APOE4(vxIs824)*; *vhl-1(ok161*) animals with or without (starting at embryos) cholesterol diet to L4 followed by picking to normal NGM and culturing at 20℃. * Indicates P < 0.05, **** indicates P < 0.0001, (n > 40 animals per condition).

Elevated temperature stress causes increased levels of ROS and HIF-1 activation in *C. elegans*^32,33^. Loss of VHL-1 leads to the stabilization of HIF-1, providing a defense mechanism against hypoxic and oxidative stress (Fig. 1c). As we previously discovered that VHL-1 inactivation mitigates the morphological degeneration of dopaminergic neurons in *C. elegans* complex I mutants^34^, we examined how a *vhl-1* deletion mutation *ok161* affected the mortality of *APOE4*-transgenic *C. elegans* under 28 °C. We found that *vhl-1* deletion abolished the effect of *APOE4* on increased mortality under 28 °C, and extended lifespan in wild-type animals under 28 °C (Fig. 1d). These results establish a *C. elegans* model for rapid APOE4-induced mortality and identified potent mortality-suppressing effects of *vhl-1* LOF mutations.

APOE4 represents a lipoprotein variant with a diminished capacity for lipid recycling, resulting in intracellular accumulation of cholesterol that is highly susceptible to oxidation^35–37^. Because *C. elegans* cannot synthesize cholesterol, its cholesterol levels are determined and can be controlled by its diet. We developmentally synchronized and cultured the *APOE4*-transgenic strain on culture plates deficient in exogenously added cholesterol (Extended Data Fig. 1a-1b), a procedure to reduce overall cholesterol intake during larval development^38^. Such cholesterol-reduction conditions markedly restored the lifespan of *APOE4*-transgenic animals, without affecting that of wild type (Fig. 1e) or the mortality-decreasing effect of *vhl-1* deletion (Fig. 1f). Exogenous supplementation with N-acetyl-cysteine (NAC), a precursor of glutathione and scavenger of ROS previously used and validated in *C. elegans*^39–42^, dose-dependently suppressed the mortality effect of APOE4 (Fig. 1g). We also observed that body size was reduced in *APOE4*-transgenic *C. elegans* when compared to wild type at normal 20 °C, while *vhl-1* deletion LOF mutation or reduction of cholesterol uptake starting at embryonic stages were sufficient to rescue body sizes (Extended Data Fig. 1c-1d).

We next tested how APOE4 may interact with other genetic and environmental factors on mortality. Given that APOE4 can affect clearance of the amyloid precursor protein APP, which is also implicated in AD, we examined the neuronal *APP*-transgenic *C. elegans* under 28 °C, and found that neuronal *APP* expression did not affect mortality rates in *C. elegans* (Extended Data Fig. 1e-1f). We also observed that neuronal *APOE3* expression did not affect mortality rates under 28 °C (Extended Data Fig. 1g).

Additionally, we used a heat-independent approach to generate excessive oxidative stress based on a transgenic strain with blue light-induced production of superoxide from neuronal expression of a genetically-encoded miniSOG transgene^43,44^. We observed that blue light exposure in this strain induced a rapid and robust increase of population mortality that was strongly suppressed by dietary cholesterol reduction or NAC supplementation (Fig. 1i). *vhl-1* deletion recapitulated such mortality-suppressing effects (Fig. 1j). Furthermore, we found that *APOE4* also increased the mortality of *C. elegans* under 20 °C normal culture conditions and *vhl-1* deletion or cholesterol reduction strongly suppressed the mortality effect of *APOE4* (Fig. 1k-n and Extended Data Fig. 1h).

Taken together, these results identify VHL-1 as a potent modifier of APOE4 in mortality and suggest that APOE4 may increase intracellular cholesterol, oxidation of which by ROS contributes to an increase in population mortality suppressible by *vhl-1* deletion.

### Roles of HIF-1 in suppressing mortality caused by APOE4

We next examined roles of HIF-1 in suppressing mortality. We monitored hypoxic and redox stress responses using the well-characterized HIF-1-dependent transcriptional reporter, *cysl-2*p::GFP^45–47^. As would be predicted for stabilized HIF-1, *vhl-1* deletion strongly activated *cysl-2*p::GFP in a HIF-dependent manner (Fig. 2b and Extended Data Fig. 2a). Under normal 21% oxygen conditions, elevated temperature at 28 °C caused a time- and temperature-dependent activation of *cysl-2*p::GFP (Extended Data Fig. 2b-2d), consistent with elevated oxidative stress and HIF-1 activation by heat^32^. LOF *hif-1* fully suppressed the mortality-reducing effects of *vhl-1* under both normal culture conditions^26,48^ and on *APOE4* at 28 °C (Fig. 2a and 2c). We focused on characterizing the effects of a stabilized form of HIF-1 using a transgene *otIs197* that expresses a non-degradable (VHL-resistant) P621A variant and driven by the *unc-14* promoter^49^ (Fig. 2d). Testing thermal stress, we found that stabilized HIF-1 extended the lifespan of wild type grown at 28 °C (Fig. 2e) and suppressed the mortality effect of *APOE4* to the same level as *vhl-1* deletion (Fig. 2f). Testing cholesterol as a stressor, we found that reducing cholesterol during larval development but not during adult stage occluded negative effects of APOE4 in both wild type and stabilized HIF-1 transgenic *otIs197* animals (Fig. 2g-2i). In addition, supplementation with NAC dose-dependently reduced mortality of APOE4 but to a lesser extent in stabilized HIF-1 or *vhl-1* deletion mutant animals (Fig. 2j). Furthermore, stabilized HIF-1 also recapitulated the effect of *vhl-1* deletion on reducing the mortality of *APOE4* transgenic animals at 20 °C (Fig. 2k-2n).

**Figure 2.**
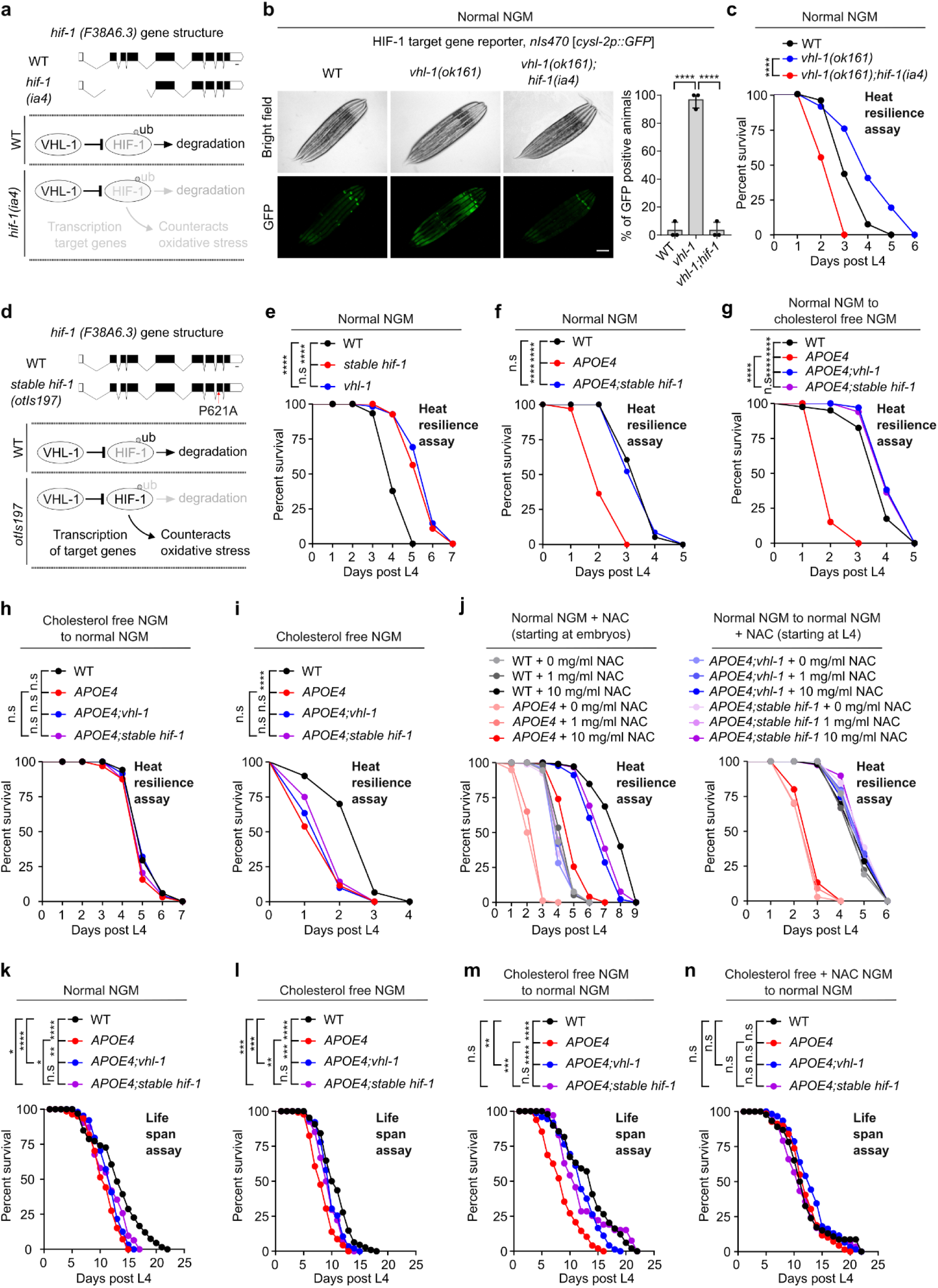
Stabilized HIF-1 recapitulates the effects of VHL-1 inactivation. (a) Schematic of the *hif-1(ia4)* LOF deletion allele (1,231 bp deletion of the second, third, and fourth exons) and its impaired capacity to counteract oxidative stress. Scale bar: 100 bp. (b) Representative epifluorescence images and quantification showing that *cysl-2p::GFP* constitutive upregulation in *vhl-1* mutants is blocked by *hif-1(ia4)*. Scale bar: 100 μm. ****indicates P < 0.0001 (n > 30 animals per condition). (c) Lifespan curves of WT, LOF mutant *vhl-1(ok161)* and double LOF mutant *vhl-1* (*ok161*); *hif-1(ia4)* animals at 28 ℃ starting at L4 on normal NGM. **** indicates P < 0.0001 (n > 40 animals per condition). (d) Schematic of non-degradable form of HIF-1 (P621A) expressed by the *unc-14* promoter (predominantly active in neurons) in *hif-1* mutant background (*otIs197 [unc-14p::hif-1(P621A) + ttx-3p::RFP*]. Scale bar: 100 bp. (e) Lifespan curves of WT, non-degradable form of HIF-1(P621A) (*otIs197*) or *vhl-1*(*ok161*) LOF mutant animals at 28 ℃ starting at L4 on normal NGM. **** indicates P < 0.0001, n.s indicates non-significant (n > 40 animals per condition). (f) Lifespan curves of WT, *APOE4(vxIs824); otIs197* and *APOE4(vxIs824)* animals at 28 ℃ starting at L4 on normal NGM. **** indicates P < 0.0001, n.s indicates non-significant (n > 40 animals per condition). (g) Lifespan curves of WT, *APOE4(vxIs824), APOE4(vxIs824)*; *vhl-1(ok161*) and *APOE4(vxIs824); otIs197* animals grown to L4 on normal NGM followed by picking to cholesterol free NGM and culturing at 28℃. *** indicates P < 0.001, n.s indicates non-significant (n > 40 animals per condition). (h) Lifespan curves of WT, *APOE4(vxIs824)*, *APOE4(vxIs824)*; *vhl-1(ok161*) and *APOE4(vxIs824); otIs197* animals starting at early life (embryos) with cholesterol free NGM to L4 followed by picking to normal NGM and culturing at 28℃. n.s indicates non-significant (n > 40 animals per condition). (i) Lifespan curves of WT, *APOE4(vxIs824), APOE4(vxIs824)*; *vhl-1(ok161*) and *APOE4(vxIs824); stable hif-1* (*otIs197*) animals starting at early life (embryos) with cholesterol free NGM to L4 followed by picking to cholesterol free NGM and culturing at 28℃. **** indicates P < 0.0001, n.s indicates non-significant, (n > 40 animals per condition). (j) Lifespan curves of WT, *APOE4(vxIs824)*, *APOE4(vxIs824)*; *vhl-1(ok161*) and *APOE4(vxIs824); stable hif-1* (*otIs197*) animals starting at early life (starting at embryos) with indicated NAC concentration diet to L4 on normal NGM followed by picking to normal NGM supplemented with indicated concentration of NAC and transferred to 28℃ (left). Lifespan curves of WT, *APOE4(vxIs824)* and *APOE4(vxIs824); stable hif-1* (*otIs197*) animals grown to L4 on normal NGM followed by picking to normal NGM supplemented with indicated concentration of NAC (starting at L4 stage) and culturing at 28℃. (n > 40 animals per condition). (k) Lifespan curves of WT, *APOE4(vxIs824)*, *APOE4(vxIs824)*; *vhl-1(ok161*) and *APOE4(vxIs824); stable hif-1* (*otIs197*) animals at constant 20 ℃ on normal NGM. * Indicates P < 0.05, ** indicates P < 0.01, *** indicates P < 0.001, **** indicates P < 0.0001, n.s indicates non-significant (n > 40 animals per condition). (l) Lifespan curves of WT, *APOE4(vxIs824), APOE4(vxIs824)*; *vhl-1(ok161*) and *APOE4(vxIs824); stable hif-1* (*otIs197*) animals at constant 20 ℃ on cholesterol free NGM. ** indicates P < 0.01, *** indicates P < 0.001, **** indicates P < 0.0001, n.s indicates non-significant (n > 40 animals per condition). (m) Lifespan curves of WT, *APOE4(vxIs824), APOE4(vxIs824)*; *vhl-1(ok161*) and *APOE4(vxIs824); stable hif-1* (*otIs197*) animals starting at early life (start at embryos) with cholesterol free NGM to L4 followed by picking to normal NGM and culturing at 20℃. ** indicates P < 0.01, *** indicates P < 0.001, **** indicates P < 0.0001, n.s indicates non-significant (n > 40 animals per condition). (n) Lifespan curves of WT, *APOE4(vxIs824)*, *APOE4(vxIs824); vhl-1(ok161)* and *APOE4(vxIs824); stable hif-1 (otIs197)* animals starting at early life (embryos) with cholesterol free and 10 mg/ml NAC diets to L4 on cholesterol free NGM followed by picking to normal NGM and culturing at 20℃ of incubator. n.s indicates non-significant (n > 40 animals per condition).

To test if HIF-1 played a similar role beyond *C. elegans*, we also generated a HEK293T cell line by expressing stabilized mammalian HIF-1 by lentiviral infection (Extended Data Fig. 2e). We found that it similarly protected HEK293T cells against thermal stress conditions and suppressed the mortality-increasing effect of APOE4 (Extended Data Fig. 2f). The abundance, subcellular localization and secretion of APOE4 were not apparently affected by stabilized HIF-1 or thermal stress in HEK293T cells (Extended Data Fig. 2g-2h). Exogenous supplementation with *APOE4*-expressing HEK293T cell supernatants did not affect the mortality of *C. elegans* under 28 °C (Extended Data Fig. 2i). In HEK293T cells, we found that stabilized HIF-1 can also suppress heat or *APOE4*-induced genomic DNA fragmentation (Extended Data Fig. 2j).

Together, these results demonstrate roles of HIF-1 in mediating effects of *vhl-1* loss in mortality and that a stabilized HIF-1 transgene is sufficient to suppress APOE4-induced increase in mortality during normal aging and under heightened heat stress conditions.

### Cellular consequences of APOE4 suppressed by *vhl-1* loss or HIF-1 activation

To understand mechanisms of APOE4 toxicities and protection by *vhl-1* and HIF-1, we assessed the molecular and cellular abnormalities in neuronal *APOE4* transgenic animals. To discover pathways potentially dysregulated by APOE4, we performed transcriptome profiling. RNAseq analysis revealed that APOE4 caused numerous alterations in genes involved in stress responses and proteostasis (Fig. 3a). To monitor proteostasis *in vivo*, we generated a transcriptional reporter for the heat shock protein-encoding *hsp-16.2* as a live indicator. We found that *hsp-16.2*p::GFP remained at low baseline levels throughout development in the wild type under normal culture conditions (Fig. 3b and Extended Data Fig. 3a). By comparison, APOE4 increased *hsp-16.2*p::GFP expression dramatically starting at fourth larval stage and with the highest penetrance at day 5 post L4 (Fig. 3b and Extended Data Fig. 3c). APOE4 elevated proteostatic stress, as revealed by this reporter, even without exogenous proteostasis-perturbing conditions, such as heat stress. High-magnification confocal microscopic analysis revealed the site of abnormally up-regulated *hsp-16.2*p::GFP expression predominantly in the body wall muscle, while its expression in a few unidentified neurons remained largely unaltered (Fig. 3d-3f). As a more direct readout of proteostasis^50^, we also monitored length-dependent aggregation of polyQ-YFP fusion proteins in *C. elegans*. We found that APOE4 increased *unc54p::Q40::YFP* aggregation, but not *unc54p::Q35::YFP* in the body wall muscle (Fig. 3f, Extended Data Figs. 3b-3c). To monitor the proteotoxic consequences of APOE4 and *vhl-1*, we used Western blot to assess oxidative stress-induced actin cleavage (Fig. 3h). While APOE4 caused dramatic accumulation of actin species with lower molecular weight indicative of protein cleavage, such proteotoxic effects were largely absent in *vhl-1* LOF deletion mutants or stabilized HIF-1 animals carrying APOE4 (Fig. 3h). Actin cleavage also occurred in wild-type animals subjected to 28 °C heat stress, and was similarly suppressed in *vhl-1* deletion mutants or stabilized HIF-1 animals without APOE4 (Fig. 3h). Immunocytochemistry showed that the antibody used for Actin stained mostly body wall muscles, consistent with *hsp-16.2*p::GFP activation in the same tissue (Fig. 3i). Given neuronal specific APOE4 expression, these results suggest non-cell autonomous proteotoxic effects of APOE4 suppressible by *vhl-1* loss or HIF-1 activation.

**Figure 3.**
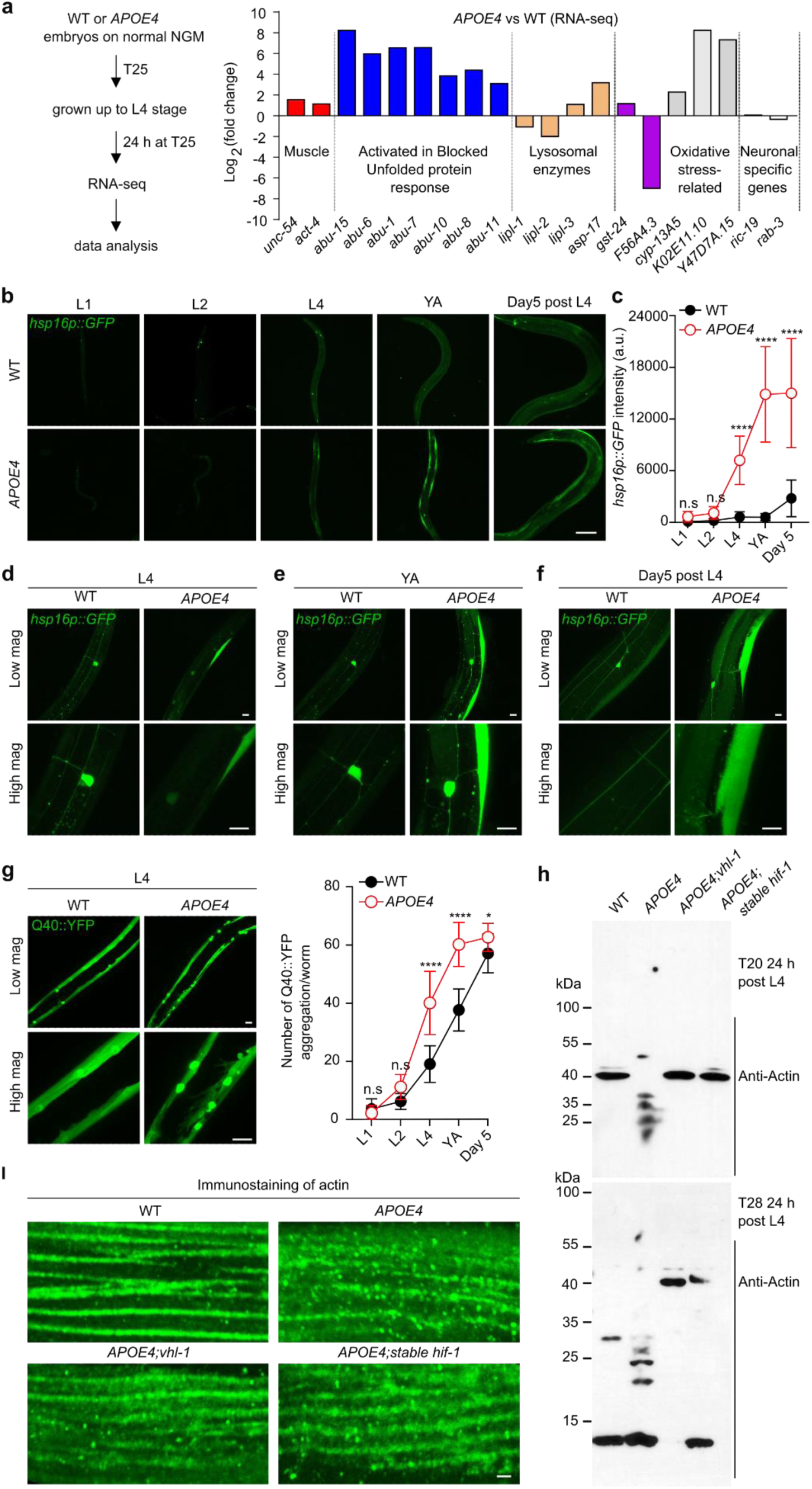
*APOE4* causes non-cell autonomous proteostasis dysregulation and actin cleavage suppressed by *vhl-1*. (a) Schematic for RNA-seq transcriptome profiling of WT and pan neuronal transgenic *APOE4 (vxIs824*), showing representative genes of various classes dysregulated in *APOE4* compared to WT. (b) Representative confocal low-magnification images of *hsp16p::GFP* in body wall muscles in WT and *APOE4 (vxIs824*) animals at different stages of L1, L2, L4, young adult (day1 post L4) and Day 5 post L4 on normal NGM. Scale bar: 100 μm. (c) Quantification of fluorescence intensities of *hsp16p::GFP* in body wall muscles under conditions indicated. *** indicates P < 0.001, n.s indicates non-significant (n > 30 animals per condition). (d-f) Representative confocal high-magnification images of *hsp16p::GFP* in body wall muscles in WT and *APOE4 (vxIs824*) at different stages of L4, young adult (day1 post L4) and Day 5 post L4 on normal NGM. Scale bar: 10 μm. (g) Representative confocal high-magnification images of *unc54p::Q40::YFP* in body wall muscles in WT and *APOE4 (vxIs824*) at stages of L4 on normal NGM, and quantification of aggregation number of *unc54p::Q40::YFP* in body wall muscles under conditions indicated. Scale bar: 10 μm. * indicates P < 0.05, **** indicates P < 0.0001, n.s indicates non-significant (n > 30 animals per condition). (h) Representative SDS-PAGE western blots of WT, *APOE4(vxIs824), APOE4(vxIs824)*; *vhl-1(ok161*) and *APOE4(vxIs824); stable hif-1* (*otIs197*). (i) Representative confocal high-magnification images in body wall muscles of WT, *APOE4(vxIs824), APOE4(vxIs824)*; *vhl-1(ok161*) and *APOE4(vxIs824); stable hif-1* (*otIs197*) animals immunostained with primary antibody against actin at young adult stages (24 hrs post L4) on normal NGM. Scale bar: 1 μm.

We next examined potential cell-autonomous consequences of APOE4 with respect to *vhl-1* and HIF in neurons. Given the dramatic morphological deterioration of the PVD neuron in APOE4 animals (Fig. 1b, Extended Data Figs. 4a-4d), we focused on a detailed longitudinal analysis of PVD morphological integrity in both *APOE4* and *APOE4*; *vhl-1* animals. Confocal imaging analysis revealed that the morphological defect, including decreased dendrite numbers and complexity, of the PVD neuron manifested early at the larval L4 stage and persisted throughout adulthood (Fig. 4a-4c). We found that *vhl-1* deletion strongly suppressed the morphological defects of the PVD neuron in neuronal APOE4 transgenic animals (Fig. 4a-4c). While APOE4 caused a nearly fully penetrant defect of PVD neurons at the larval L4 stage, *vhl-1* mutants exhibited marked suppression of defects in all three stages examined (Fig. 4d-4f).

**Figure 4.**
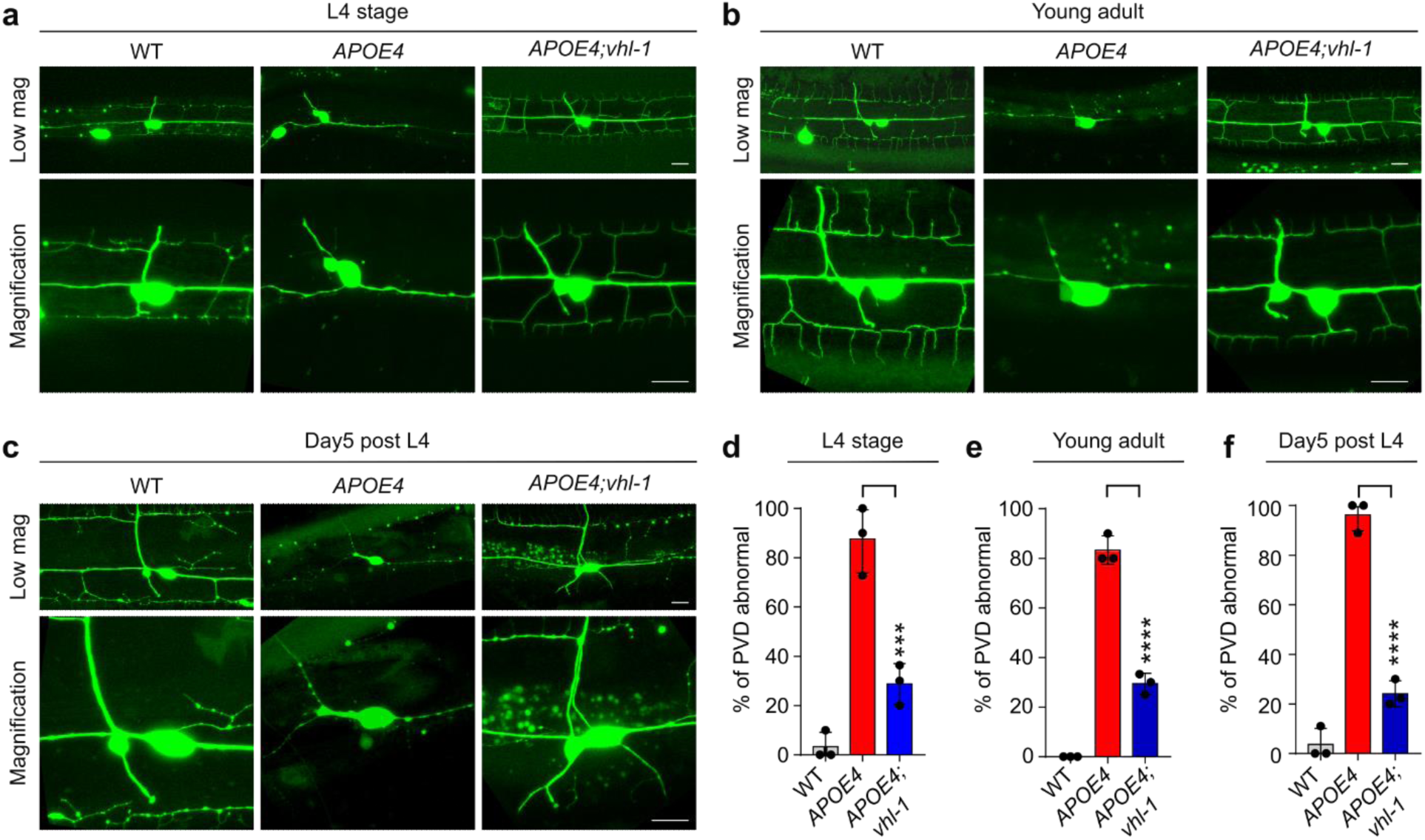
*APOE4* causes PVD morphological deterioration suppressed by *vhl-1*. (a) Representative confocal images of PVD neuron in WT, *APOE4(vxIs824), APOE4(vxIs824)*; *vhl-1(ok161*) at L4 stages on normal NGM showing *vhl-1* LOF mutants with rescued *APOE4*-induced PVD morphological deterioration. Scale bar: 10 μm. (b) Representative confocal images of PVD neuron in WT, *APOE4(vxIs824), APOE4(vxIs824)*; *vhl-1(ok161*) at young adult stages on normal NGM showing *vhl-1* LOF mutants with rescued *APOE4*-induced PVD morphological deterioration. Scale bar: 10 μm. (c) Representative confocal images of PVD neuron in WT, *APOE4(vxIs824), APOE4(vxIs824)*; *vhl-1(ok161*) at day 5 post L4 stages on normal NGM showing *vhl-1* LOF mutants with rescued *APOE4*-induced PVD morphological deterioration. Scale bar: 10 μm. (d-f) Quantification of % of PVD neuron abnormal (with the third and fourth branches of PVD neurons missing or severed) in WT, *APOE4(vxIs824), APOE4(vxIs824)*; *vhl-1(ok161*) under conditions indicated on normal NGM. *** indicates P < 0.001, (n > 30 animals per condition).

Together, these results show that APOE4 can cause both non cell-autonomous and cell-autonomous cellular defects, both of which are suppressible by *vhl-1* LOF. To further investigate cellular mechanisms underlying the neuronal toxicity of APOE4 and protection by *vhl-1* or HIF-1, we examined major organelles in live neurons, including mitochondria, endosomes, and lysosomes. Using the neuronal organelle-specific fluorescent markers (schematic in Fig. 5a) for longitudinal imaging, we found that APOE4 did not appear to affect neuronal endosomes, as compared to wild type (Extended Data Fig. 5a). However, APOE4 caused a striking age-dependent increase of the fluorescent marker for mitochondria (Fig. 5b-5e) and decrease of the fluorescent marker for lysosomes (Fig. 5h). The increase of mitochondrial markers did not manifest until fourth larval stage and persisted throughout adult stage (Fig.5c). The changes in organelle reporters could not be explained by APOE4 affecting transgene expression since RNAseq results (Table S1, Fig. 3a) indicated that APOE4 does not affect the expression of *ric-19*, the promoter of which drives the organelle markers. Strikingly, *vhl-1* deletion or stabilized HIF-1 strongly suppressed the abnormally increased mitochondrial markers by APOE4 (Fig. 5f-5g). Reduction of cholesterol also suppressed the effect of APOE4 on such mitochondrial and lysosomal phenotypes (Extended Data Fig. 5c-5e). These results reveal organelle-specific defects caused by APOE4 and suggest that APOE4 likely exerts cellular toxicity through excess cholesterol, oxidation of which leads to lysosomal membrane disruption, impaired mitophagy and mitochondria clearance, defects suppressible by *vhl-1* inhibition and HIF-1 activation.

**Figure 5.**
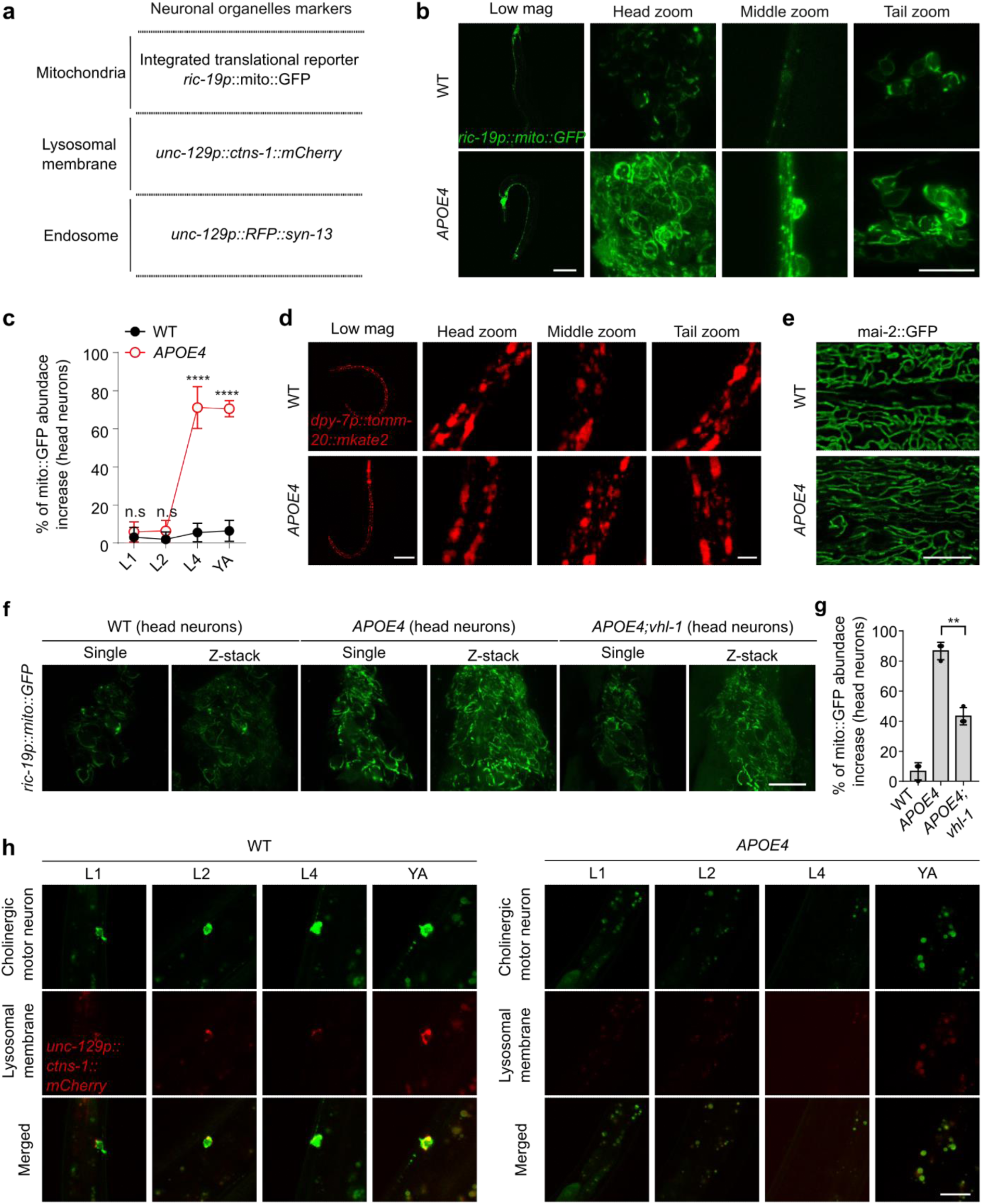
*APOE4* causes neuronal mitochondria defects suppressed by *vhl-1*. (a) Schematic of neuronal organelle-specific fluorescent markers. (b) Representative confocal low and high magnification images of neuronal tissue specific expression mitochondria reporter (*ric19p::mito::GFP*) in WT and *APOE4(vxIs824)* animals at young adult (day1 post L4 stages) with indicated position. Scale bar: 100 μm (low magnification) and 10 μm (high magnification). (c) Quantification of percentage of *ric19p::mito::GFP* abnormal based on head neurons in WT and *APOE4(vxIs824)* animals at different stages of L1, L2, L4 and young adult (day1 post L4 stages) on normal NGM. **** indicates P < 0.0001, n.s indicates non-significant (n > 30 animals per condition). (d) Representative confocal low and high magnification images of hypodermal cell mitochondria based on *dpy7p::mito::mkate2* showing no apparent change in WT and *APOE4(vxIs824)* at young adult stages on normal NGM. Scale bars: 100 μm (low magnification) and 10 μm (high magnification). (e) Representative confocal images of intestinal mitochondria based on *mai-2*::GFP showing no apparent change in WT and *APOE4(vxIs824)* at young adult stages on normal NGM. Scale bar: 10 μm. (f) Representative confocal images of *ric19p::mito::GFP* in WT, *APOE4(vxIs824)* and *APOE4(vxIs824)*;*vhl-1*(ok161) at young adult stages with head neuron positions (day1 post L4 stages) on normal NGM. Scale bar: 10 μm. (g) Quantification of percentage of *ric19p::mito::GFP* animals abnormal based on head neurons in WT, *APOE4(vxIs824)* and *APOE4(vxIs824)*;*vhl-1*(*ok161*) at young adult stages (day1 post L4 stages) on normal NGM. ** indicates P < 0.01 (n > 30 animals per condition). (h) Representative confocal images of neuronal lysosomal membrane reporter in WT and *APOE4(vxIs824)* at different stages of L1, L2, L4 and Day 1 post L4 on normal NGM. Scale bar: 10 μm.

### Transcriptional targets of HIF-1 mediating effects of *vhl-1* and HIF-1

We aimed to determine the transcriptional targets of HIF-1 and their mechanisms of action underlying protection against heat stress and APOE4. Proteomic and transcriptomic studies have identified many genes differentially regulated in *vhl-1* mutants^51–53^. We used quantitative RT-PCR (qRT-PCR) and GFP reporters to validate many of these targets based on their dramatic up-regulation in *vhl-1* mutants grown at 28 °C, under which condition HIF-1 is both stabilized and activated in target gene transcriptional transactivation (Fig. 6a). We used deletion mutants or RNAi (when deletion mutants were not available) against these candidate genes to test whether any are functionally important for survival (measured as median lifespan) at 28 °C in both wild type and *vhl-1* mutants. We found that genetic deletion or RNAi against each of two candidate genes, *tgn-38* and *Y70C5C.1*, led to increased mortality at 28 °C (Fig. 6b-6e). *tgn-38* encodes a *C. elegans* orthologue of human C5orf15 (chromosome 5 open reading frame 15) and TGOLN2 (trans-golgi network protein 2) with uncharacterized biological functions, whereas *Y70C5C.1* encodes a *C. elegans* ortholog of human IDE (insulin degrading enzyme). Though mechanisms linking TGN-38 to mortality regulation remain unclear, the loss-of-function phenotype of *Y70C5C.1* suggests that HIF-1 may activate expression of an insulin-degrading enzyme, leading to insulin receptor (DAF-2) inhibition and activation of the DAF-16 stress-responding pathway.

**Figure 6.**
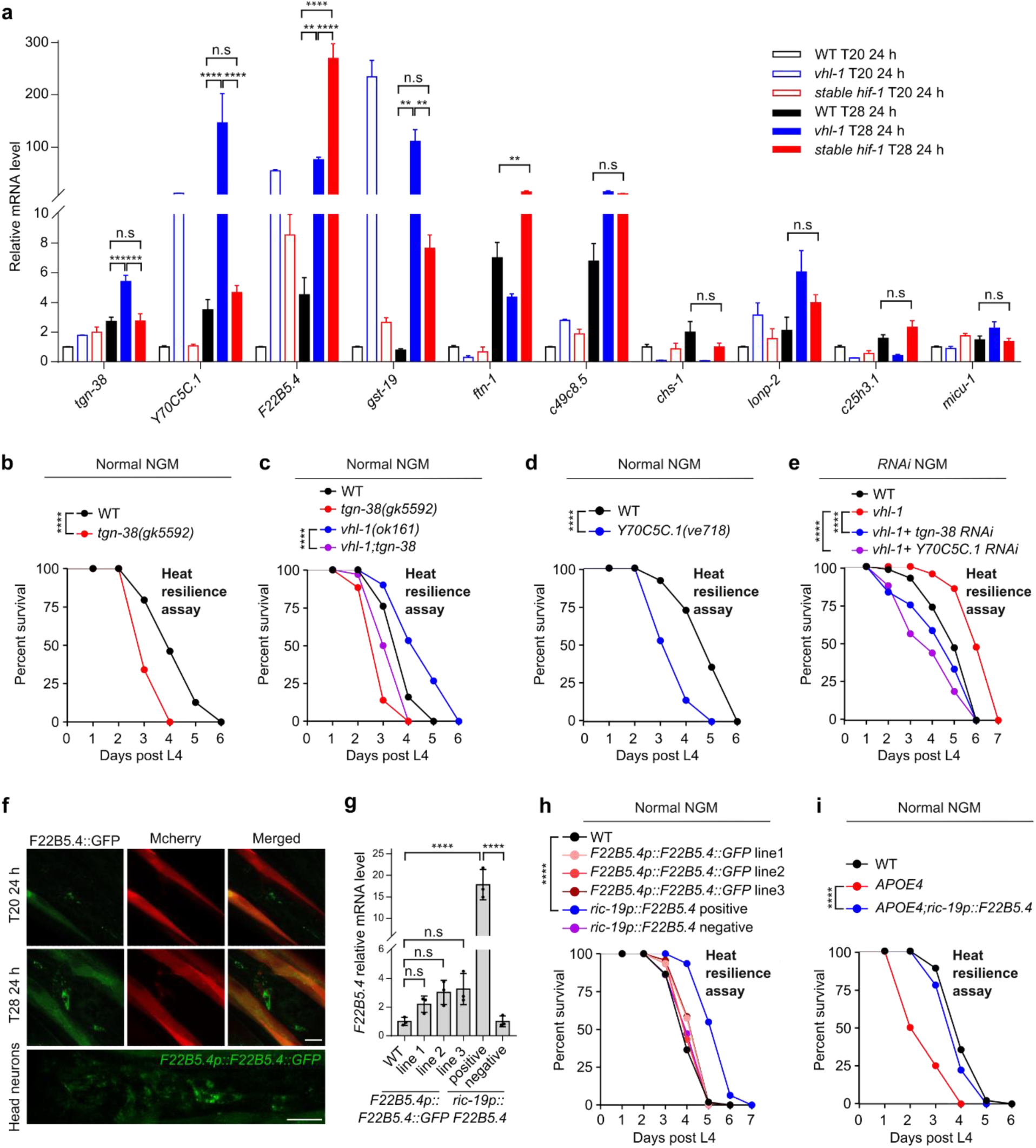
HIF-1 target gene *F22B5.4* in neurons protects against thermal stress. (a) Quantitative RT-PCR measurements of indicated gene expression levels in WT, *vhl-1(ok161*) and *otIs197* animals upon sustained treatment at 28 ℃ or 20 ℃ for 24 hours starting at L4 on normal NGM. ** indicates P < 0.01, *** indicates P < 0.001, **** indicates P < 0.0001, n.s indicates non-significant. (b-c) Lifespan curves of WT, *tgn-38(gk5592)* LOF mutants*, vhl-1(ok161)* mutants, and double LOF mutant *vhl-1; tgn-38* at 28 ℃ starting at L4 on normal NGM. **** indicates P < 0.0001 (n > 40 animals per condition). (d) Lifespan curves of WT and *Y70C5C.1(ve718)* LOF mutants at 28 ℃ starting at L4 on normal NGM. **** indicates P < 0.0001 (n > 40 animals per condition). (e) Lifespan curves of WT, *vhl-1(ok161*) mutants, *vhl-1(ok161*) mutants with RNAi against *tgn-38* and *Y70C5C.1* at 28 ℃ starting at L4. **** indicates P < 0.0001 (n > 40 animals per condition). (f) Representative confocal high magnification images of the *F22B5.4* translational reporter with GFP observed predominantly in head neurons in WT animals. Scale bar: 10 μm. (g) Quantitative RT-PCR measurements of *F22B5.4* gene expression levels under conditions indicated on normal NGM. **** indicates P < 0.0001, n.s. indicates non-significant. (h) Lifespan curves of WT, three representative F22B5.4 translational reporter lines and *ric19p::F22B5.4* over-expression gain-of-function animals at 28 ℃ starting at L4 on normal NGM. **** indicates P < 0.0001 (n > 40 animals per condition). (i) Lifespan curves of WT, *APOE4(vxIs824)* and *APOE4(vxIs824)*; Ex[*ric19p::F22B5.4*, *unc54p::mcherry*] animals at 28 ℃ starting at L4 on normal NGM. *** indicates P < 0.001 (n > 40 animals per condition).

Among the most dramatically up-regulated gene by HIF-1 (via stabilized HIF-1 or loss of *vhl-1* at 28 °C), *F22B5.4* encodes a predicted mitochondrial protein (with the probability of mitochondrial presequence of 0.967, mitoFate^54^) of uncharacterized biological function. Although we did not observe the RNAi phenotype of *F22B5.4* (likely owing to a paralogous gene *F36A2.7* and/or low RNAi efficiency in tissue of expression), single-cell gene expression profiling by CeNGEN indicates its predominant expression in neurons^55^. We generated a translational GFP reporter for *F22B5.4* under the control of its endogenous promoter and confirmed its specific expression in neurons and up-regulation by HIF-1 and in the *vhl-1* mutant (Fig. 6f). Neuronal-specific gain-of-function of *F22B5.4* by *ric-19* promoter-driven cDNA expression markedly reduced mortality at 28 °C (Fig. 6g-6h). Neuronal-specific gain-of-function of *F22B5.4* also partially suppressed the mortality phenotype caused by transgenic APOE4 (Fig. 6i).

These results identify three previously uncharacterized HIF-1 targets that functionally contribute to protection of neurons and suppression of animal mortality in *C. elegans*.

### Vhl inactivation suppresses APOE4-induced neurovascular injuries in mice

To further evaluate evolutionarily conserved mechanisms by which VHL inactivation may ameliorate toxic effects of APOE4, we assessed the neurovascular injuries in *APOE4* mice and the protective action by Vhl inhibition in mice. Human *APOE4* allele replacement in mice can lead to cerebral vascular and blood-brain barrier (BBB) lesions accompanied by compromised tight junctions, and neurodegenerative changes, including synaptic loss^36,56,57^. To investigate the potential neurovascular benefits of Vhl inactivation in such humanized *APOE4* transgenic mice, we injected AAV-*Vhl*-shRNA bilaterally into the hippocampus (Fig. 7a and Extended Data Fig. 6a). We found that the *APOE4* mice exhibited marked loss of brain capillary pericyte coverage in the hippocampus compared to the wild-type control (C57BL/6 mice). Inhibition of *Vhl* by shRNA markedly restored pericyte coverage of brain capillaries (Fig. 7b and 7c). We also observed reduced abundance of the tight junction protein, Occludin, in the brains of *APOE4* mice, which was mitigated by *Vhl* inhibition (Fig. 7d and 7e). We assessed the integrity of the BBB by intravenous injection of Evans blue dye (Fig. 7f) and found that the *APOE4* mice exhibited markedly weakened BBB as evidenced by higher optical density at 620 nm compared to C57BL/6 mice. In contrast, Evans blue content analyses showed that the BBB was largely intact when *Vhl* was knocked down in the brains of *APOE4* mice, reaching levels comparable to those observed in control C57BL/6 mice (Fig. 7f). In addition, we observed that *APOE4* caused a striking loss of hippocampal axons and decreased protein levels of the synaptic marker Synaptophysin in the brain, whereas inhibition of *Vhl* markedly reversed both axonal and synaptic degeneration phenotypes caused by *APOE4* (Fig. 7g-7j). Collectively, these findings demonstrate that genetic inhibition of *Vhl* can strongly ameliorate *APOE4*-induced cerebrovascular injuries and neuronal synaptic damage in mice.

**Figure 7.**
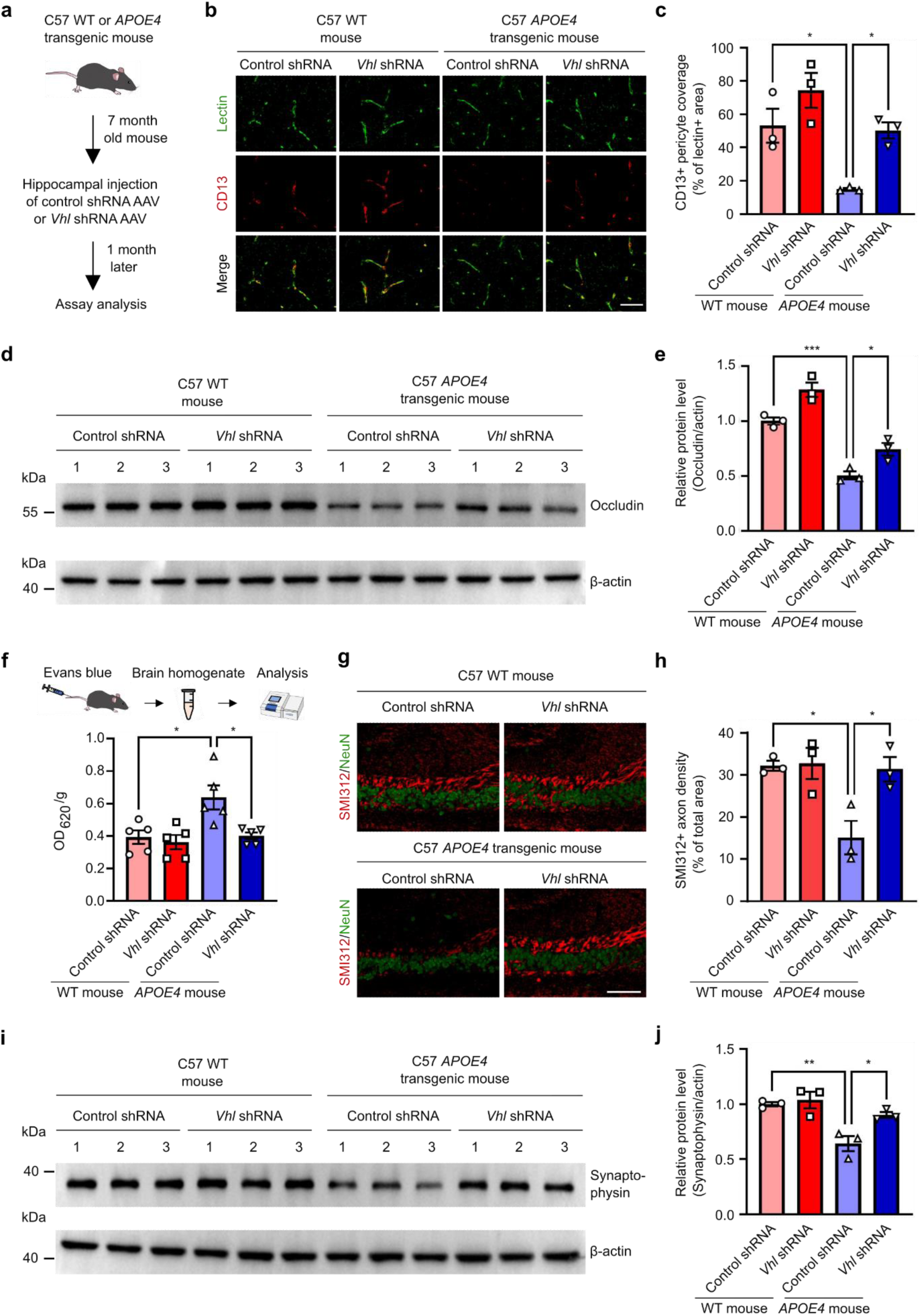
*Vhl* inhibition mitigates cerebral vascular and synaptic damages in humanized *APOE4* transgenic mice. (a) Schematic for the knockdown of *Vhl* by AAV-shRNA in humanized APOE4 transgenic mice. (b-c) Representative images of CD13+ pericyte coverage (red) of lectin+ endothelial capillary profiles (green) in the hippocampus (b). Quantification of pericyte coverage on capillaries (c). * indicates P < 0.05, n = 3 mice per group. Scale bar: 50 μm. (d-e) Representative western blot showing occludin proteins from mouse brain tissues (d) and quantification of relative protein levels of Occludin (e). * indicates P < 0.05, *** indicates P < 0.001, n = 3 mice per group. (f) Schematic for the Evans blue leakage experiment and quantification of Evans blue leakage in mouse brain tissues. * indicates P < 0.05, n = 5 mice per group. (g-h) Representative images of SMI312+ axons (red) and NeuN+ neurons (green) in the hippocampus (g), with quantification of SMI312+ axon density (h). * indicates P < 0.05, n = 3 mice per group. Scale bar: 100 μm. (i-j) Representative western blot showing Synaptophysin proteins from mouse brain tissues (i). Quantification of relative protein levels of Synaptophysin (j). * indicates P < 0.05, ** indicates P < 0.01, n = 3 mice per group. Data were presented as means ± S.E.M.

## Discussion

Age-related mortality represents a universal phenomenon influenced by intrinsic genetic factors, environmental stressors, and stochastic events. In this study, we investigated how the VHL-HIF axis modulates mortality and cell damage in *C. elegans* and mice. Our findings reveal that targeting VHL-1 remarkably suppresses mortality induced by various factors, including elevated ROS, temperature stress, and the expression of the human *APOE4* gene variant associated with neurodegeneration and mortality in humans. We established a *C. elegans* model for rapid APOE4-induced mortality and demonstrated the mortality-suppressing effects of VHL-1 inactivation. We show that stabilized HIF-1 recapitulates the effects of VHL-1 inactivation, likely through orchestrating a genetic program that defends against various cellular dysfunctions linked to mortality, including mitochondrial abnormalities, oxidative stress, proteostasis dysregulation, and endo-lysosomal rupture (Extended Data Fig. 7). We identified previously uncharacterized genes, including *tgn-38, Y70C5C.1*, and *F22B5.4*, as HIF-1 targets that contribute to mortality suppression, adding depth to a molecular mechanistic understanding.

Extensive studies have investigated mechanisms of cellular toxicity associated with APOE4 in the context of neurodegeneration and AD. Emerging evidence suggests that neuronal APOE4 may act as a crucial upstream trigger and likely a driver of late-onset AD pathogenesis, leading to downstream neuro-inflammation, glial responses and subsequent neurodegeneration^58^. Our study sheds light on the cellular consequences of neuronal APOE4 expression, revealing not only cell-autonomous effects of APOE4 in promoting neuronal morphological deterioration, mitochondrial dysfunction and lysosomal disruption in neurons, but also cross-tissue actions on proteostatic abnormalities in body wall muscles. Neuronal APOE4 inflicts oxidative stress via excess ROS generation and intracellular cholesterol accumulation by multiple mechanisms^58–60^, which may separately and additively lead to the observed cellular defects in *C. elegans*. Importantly, reduction of cholesterol from dietary sources or amelioration of excess oxidative stress through NAC or HIF-1 stabilization strongly suppressed these defects, providing a causal link from cholesterol to mortality regulation by VHL-HIF.

In mice, we showed that *Vhl* knockdown mitigated neurovascular injuries induced by APOE4. Beneficial effects of targeting Vhl in neural tissues include enhanced pericyte coverage, preservation of tight junction proteins, and protection against blood-brain barrier compromise and synaptic loss. This evidence of a conserved mechanism in a mammalian system strengthens the potential clinical implications of targeting VHL-HIF for mitigating age-related mortality and neurodegenerative risks associated with APOE4. Although *Vhl* loss or HIF-1 activation in dividing cells could be oncogenic, leading to tumor cell growth, specific targeting of VHL-HIF in non-proliferative tissues, such as post-mitotic neurons, might broadly protect against oxidative stress resulting from ischemia-reperfusion injuries, neurodegeneration, aging or *APOE4* genetic predisposition. The integration of our findings across different species paves the way for future studies into conserved mechanistic links underlying the complex relationships among genetic factors, cellular pathways, and environmental influences on mortality.

The reconceptualization of APOE4 homozygosity as a form of genetically determined AD highlights the importance of understanding APOE4 biology, pathophysiology and identifying targets that can be harnessed to modify APOE4-induced pathologies in cells and organisms. We use genetic, cell biological and phenotypic analyses to elucidate how APOE4 causes toxicity in *C. elegans* and identify the VHL-HIF pathway as a potent APOE4 toxicity modifier. Our studies also raise many interesting questions that remain unanswered. The mechanisms by which the three HIF-1 targets protect against cellular damage and animal mortality in *C. elegans* await further studies. Whole-animal genetic LOF of *vhl-1* and constitutive expression of stabilized HIF-1 in our study preclude high-resolution dissection of the spatiotemporal requirement of VHL-HIF signaling in protection against cellular damages and animal mortality. The proteostasis defects in body wall muscles and morphological deterioration of PVD neurons caused by pan-neuronal expression of *APOE4* raise intriguing cell biological questions regarding mechanisms of cross-tissue interactions, but the relative contribution of cell autonomous and non-cell autonomous effects of APOE4 to mortality in *C. elegans* remain undetermined. Although loss of *vhl-1* or HIF-1 activation protects against mortality in *C. elegans*, it remains unclear whether it is also true in mice or humans. In addition, the broader implications of VHL-HIF modulation on other aspects of organismal health, such as neurological and behavioral outcomes, warrant further investigations.

## Extended Data Figures

**Extended Data Fig. 1.**
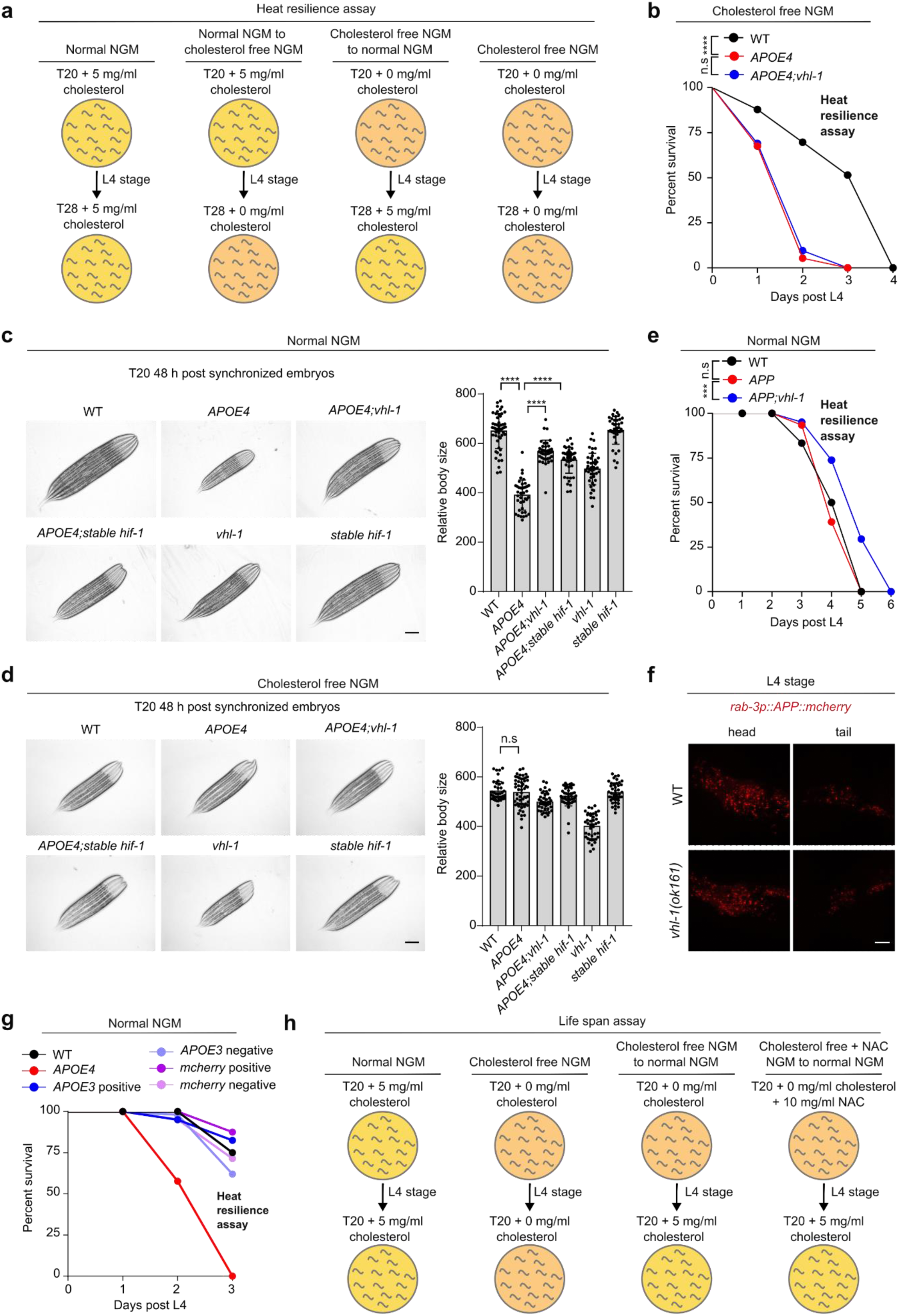
Characterization of pan-neuronal expression *APOE4* and the modification by *vhl-1* LOF or HIF-1 stabilization in *C. elegans*. (a) Schematic of experimental flow of animals grown to L4 stage on normal NGM followed by transfer to normal NGM for survival (at 28 ℃) assay; schematic of experimental flow of animals grown to L4 on normal NGM followed by transfer to cholesterol free NGM for survival (at 28 ℃) assay; schematic of experimental flow of animals grown to L4 on cholesterol free NGM followed by transfer to normal NGM for survival (at 28 ℃) assay; schematic of experimental flow of animals grown to L4 on cholesterol free NGM followed by transfer to cholesterol free NGM for survival (at 28 ℃) assay. (b) Lifespan curves of indicated animals at 28 ℃ starting at L4 on cholesterol free NGM. *** indicates P < 0.001, n.s indicates non-significant (n > 40 animals per condition). (c) Representative bright-field images and quantification of body size of animals with indicated genotype on normal NGM. Scale bar: 100 μm. **** indicates P < 0.001 (n > 30 animals per condition). (d) Representative bright-field images and quantification body size of animals with indicated genotype on cholesterol free NGM starting at eggs. Scale bar: 100 μm. n.s indicates non-significant(n > 30 animals per condition). (e) Lifespan curves of wild type, pan neuronal expression *APP (vxIs823 [rab-3p::APP::mCherry::unc-54 3’UTR] II.)*, *vhl-1*;*vxIs823* at 28 ℃ starting at L4 on normal NGM. *** indicates P < 0.001, n.s. indicates non-significant (n > 40 animals per condition). (f) Representative confocal images of protein APP::mcherry in head and tail neuron in *vxIs823* and *vhl-1*; *vxIs823* at L4 stage on normal NGM. Scale bar: 10 μm. (g) Lifespan curves of wild type, pan neuronal expression *APOE4* (*vxIs824*), pan neuronal extra chromosome expression of *APOE3*, *Ex[rab-3p::APOE3,myo3p::mcherry]* positive, *Ex[rab-3p::APOE3,myo3p::mcherry]* negative, *Ex[myo-3::mcherry]* positive and *Ex[myo-3::mcherry]* negative animals at 28 ℃ starting at L4 on normal NGM, (n > 40 animals per condition). (h) Schematic of experimental flow of animals grown to L4 stage on normal NGM followed by transfer to normal NGM for life span (at 20 ℃) assay; schematic of experimental flow animals grown to L4 on cholesterol free NGM followed by transfer to cholesterol free NGM for life span (at 20 ℃) assay; schematic of experimental flow of animals grown to L4 on cholesterol free NGM followed by transfer to normal NGM for life span (at 20 ℃) assay, schematic of experimental flow of animals grown to L4 on cholesterol free and supplementation with 10mg/ml NAC NGM followed by transfer to normal NGM for life span (at 20 ℃) assay.

**Extended Data Fig. 2.**
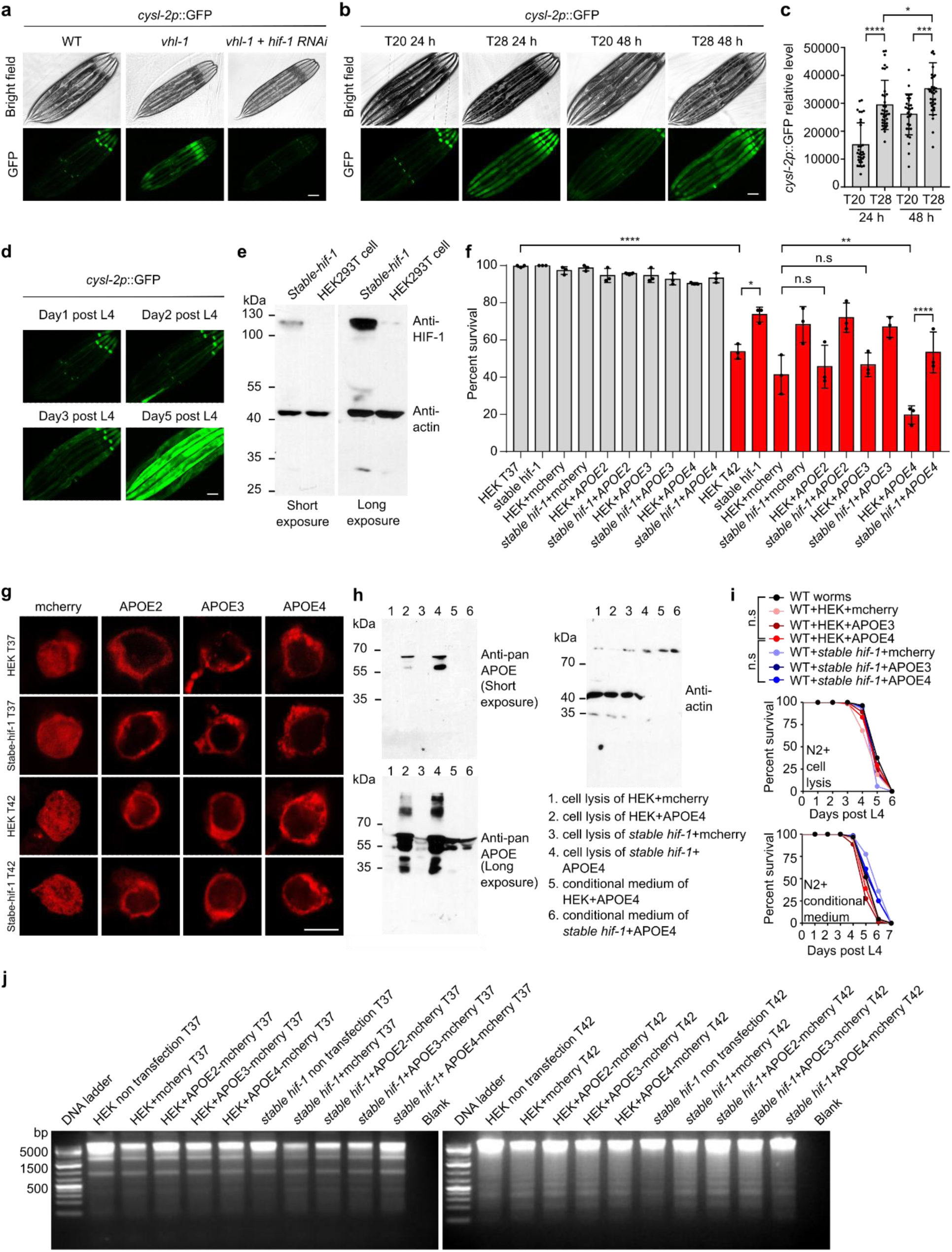
Further characterization of VHL-1, HIF-1 and APOE4. (a) Representative epifluorescence images of young adults animals showing *cysl-2p*::*GFP* constitutive upregulation in *vhl-1* mutants that can be suppressed by *hif-1* RNAi. Scale bar: 100 μm (b) Representative epifluorescence images showing *cysl-2p*::*GFP* upregulation after exposure for 24 hours or 48 hours at 28℃ starting at L4 on normal NGM. Scale bar: 100 μm (c) Quantification of fluorescence intensities of *cysl-2p*::GFP under conditions indicated. * indicates P < 0.05, *** indicates P < 0.001, **** indicates P < 0.0001, n.s indicates non-significant (n > 30 animals per condition). (d) Representative epifluorescence images showing *cysl-2p*::*GFP* upregulation during aging on normal NGM. Scale bar: 100 μm (e) Representative SDS-PAGE western blots for lysates of HEK293T cells and stable-HIF-1 HEK293T cell lines with primary antibodies against HIF-1 and actin. (f) Quantification of survival rates after heat shock in HEK293T cells showing enhanced thermal resilience conferred by gain-of-function stable-HIF-1. (g) Representative confocal images of APOE2::mCherry, APOE3::mCherry, APOE4::mCherry localization in HEK293T and stable-HIF-1 HEK293T cell line upon sustained treatment at 42 ℃ for 8 hrs. (h) Representative SDS-PAGE western blots for lysates of HEK293T cells and stable-HIF-1 HEK293T cell lines and conditional medium under indicated conditions with primary antibodies against pan-APOE and actin. (i) Lifespan curves of wild-type animals grown to L4 (starting at embryos) on normal NGM with indicated cell supernatant followed by transfer to 28 ℃ (top) and lifespan curves of wild type animals grown to L4 (starting at embryos) on normal NGM with conditional medium followed by transfer to 28 ℃ (bottom). (j) Representative agarose gel images of genomic DNA with indicated conditions.

**Extended Data Fig. 3.**
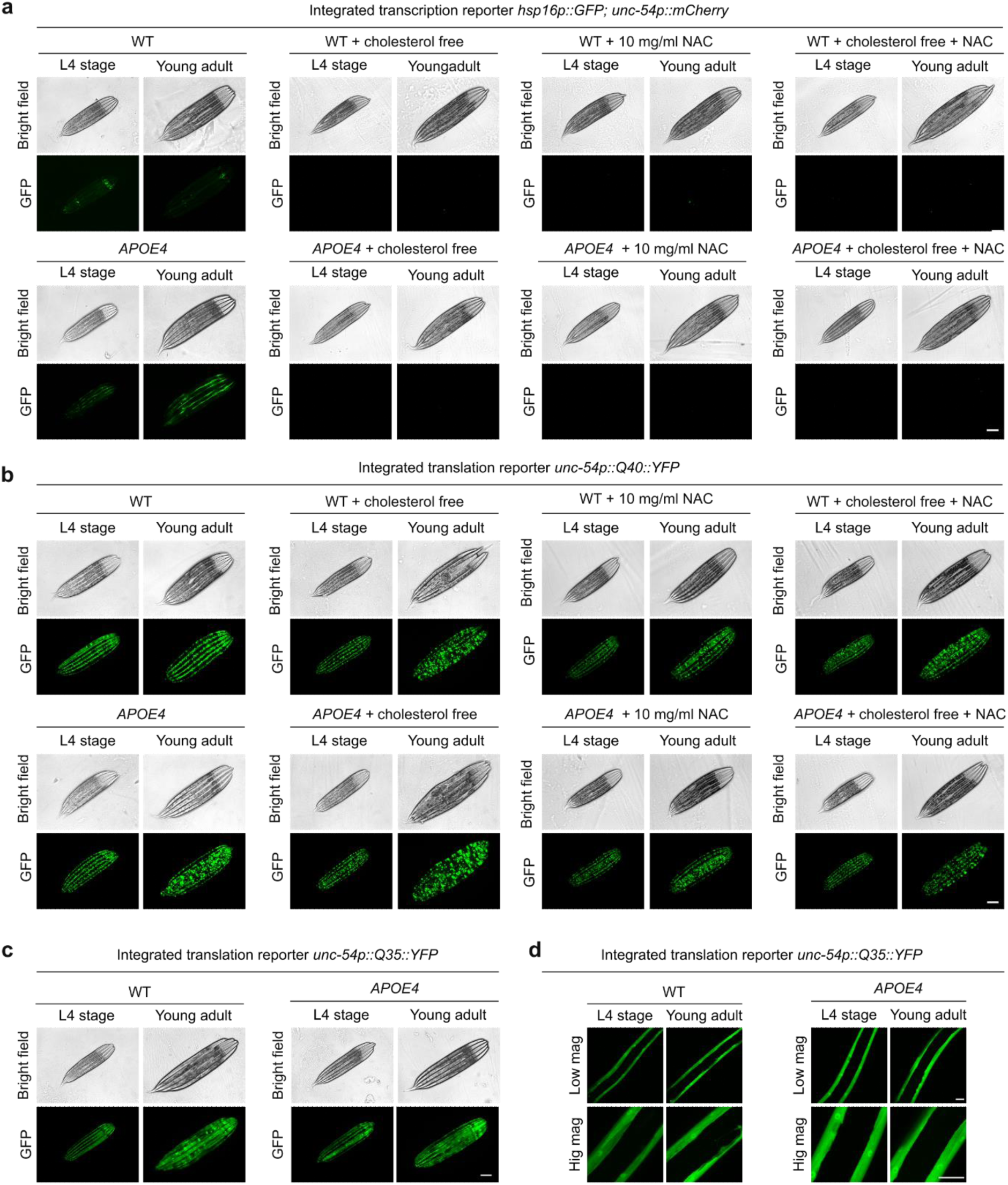
APOE4 causes proteostasis dysregulation in body wall muscles. (a) Representative bright-field and epifluorescence images of *hsp16p::GFP* in body wall muscles in WT and pan neuronal expression *APOE4(vxIs824)* on normal NGM, on cholesterol free NGM on cholesterol free NGM (starting at embryos), on normal NGM supplemented with 10 mg/ml NAC (starting at embryos), on cholesterol free NGM on cholesterol free NGM and supplemented with 10 mg/ml NAC (starting at embryos) to L4 and young adult stages. Scale bar: 100 μm. (b) Representative bright-field and epifluorescence images of *unc54p::Q40::YFP* in body wall muscles in WT and pan neuronal expression *APOE4(vxIs824)* on normal NGM, on cholesterol free NGM on cholesterol free NGM (starting at embryos), on normal NGM supplemented with 10 mg/ml NAC (starting at embryos), on cholesterol free NGM on cholesterol free NGM and supplemented with 10 mg/ml NAC (starting at embryos) to L4 and young adult stages. Scale bar: 100 μm. (c-d) Representative bright-field, epifluorescence images (c) and confocal images (d) of *unc54p::Q35::YFP* in body wall muscles in WT and pan neuronal expression *APOE4(vxIs824)* on normal NGM to L4 and young adult stages. Scale bar: 100 μm(c) and 10 μm(d).

**Extended Data Fig.4.**
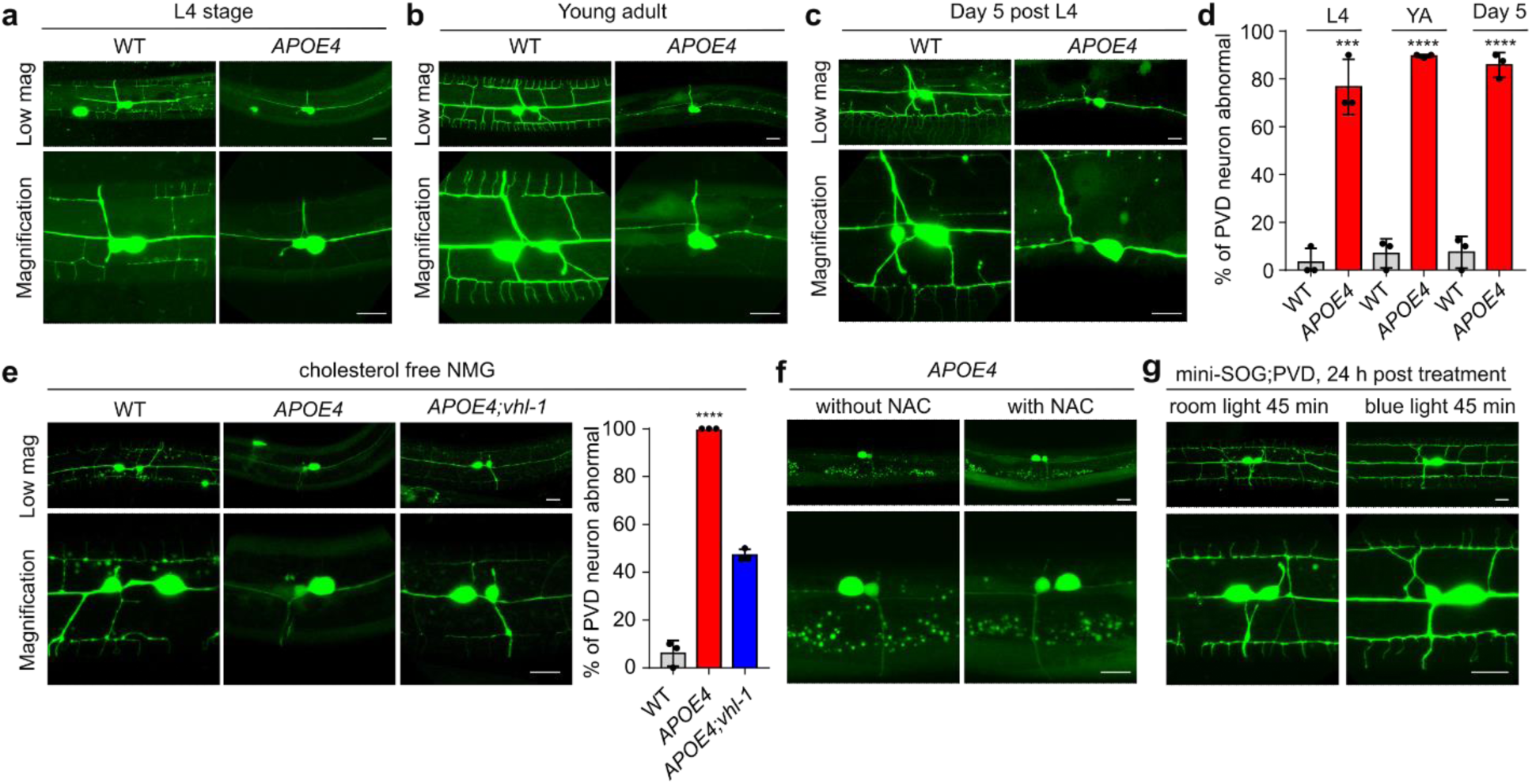
Characterization of *APOE4*-induced PVD defects. (a) Representative confocal images of PVD neuron in WT and pan neuronal expression *APOE4(vxIs824)* at L4 stages on normal NGM. Scale bar: 10 μm. (b) Representative confocal images of PVD neuron in WT and pan neuronal expression *APOE4(vxIs824)* at young adult stages (24 hrs post L4) on normal NGM. Scale bar: 10 μm. (c) Representative confocal images of PVD neuron in WT and pan neuronal expression *APOE4(vxIs824)* at day 5 post L4 stages on normal NGM. Scale bar: 10 μm. (d) Quantification of the percentage of PVD neuron abnormal (defined as the third and fourth branches of PVD neuron missing) in WT and pan neuronal expression *APOE4(vxIs824)* under conditions indicated on normal NGM. *** indicates P < 0.001, **** indicates P < 0.0001 (n > 30 animals per condition). (e) Representative confocal images and quantification of abnormal PVD neuron in WT and pan neuronal expression *APOE4(vxIs824)*, *APOE4; vhl-1* at young adult stages on cholesterol free NGM starting at embryos. Scale bar: 10 μm. (f) Representative confocal images and quantification of abnormal PVD neurons with pan neuronal expression *APOE4(vxIs824)* at young adult stages on NGM supplementation without or with 10 mg/ml NAC starting at embryos. Scale bar: 10 μm. (g) Representative confocal images of *miniSOG [unc-25p::tomm20::miniSOG::SL2::RFP]*;PVD grown to L4 on normal NGM, followed by room light and blue light treatments for 45 mins. Scale bar: 10 μm.

**Extended Data Fig. 5.**
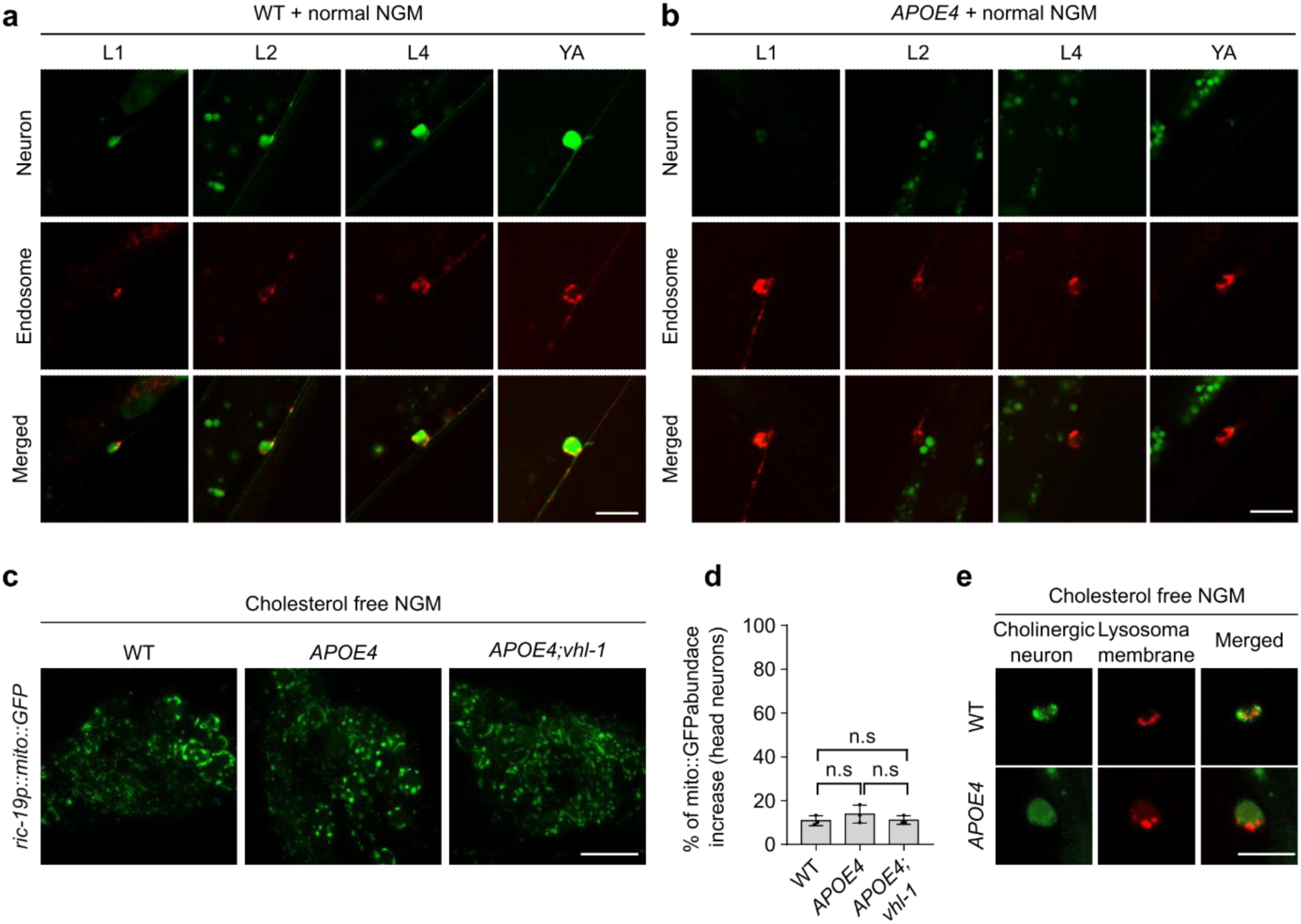
APOE4 causes abnormal neuronal mitochondria and endo-lysosomes that can be rescued by cholesterol reduction. (a-b) Representative confocal high-magnification images of neuronal endosomes in WT, pan neuronal expression *APOE4(vxIs824)* at different stages of L1, L2, L4 and YA (Day 1 post L4) on normal NGM. Scale bar: 10 μm. (c) Representative confocal high magnification images of *ric19p::mito::GFP* in WT, *APOE4(vxIs824)* and *APOE4(vxIs824)*;*vhl-1*(*ok161*) at young adults head neuron positions (day 1 post L4 stages) on cholesterol free NGM starting at embryos. Scale bar: 10 μm. (d) Quantification of the percentage of *ric19p::mito::GFP* abundance abnormal animals based on head neurons in WT, *APOE4(vxIs824)* and *APOE4(vxIs824)*;*vhl-1*(*ok161*) at young adults (day 1 post L4 stages) on cholesterol free NGM starting at embryos. n.s indicates non-significant (n > 30 animals per condition). (e) Representative confocal high-magnification images of neuronal lysosomal membrane reporter in WT and *APOE4(vxIs824)* at young adults (day 1 post L4 stages) on cholesterol free NGM starting at embryos. Scale bar: 10 μm.

**Extended Data Fig. 6.**
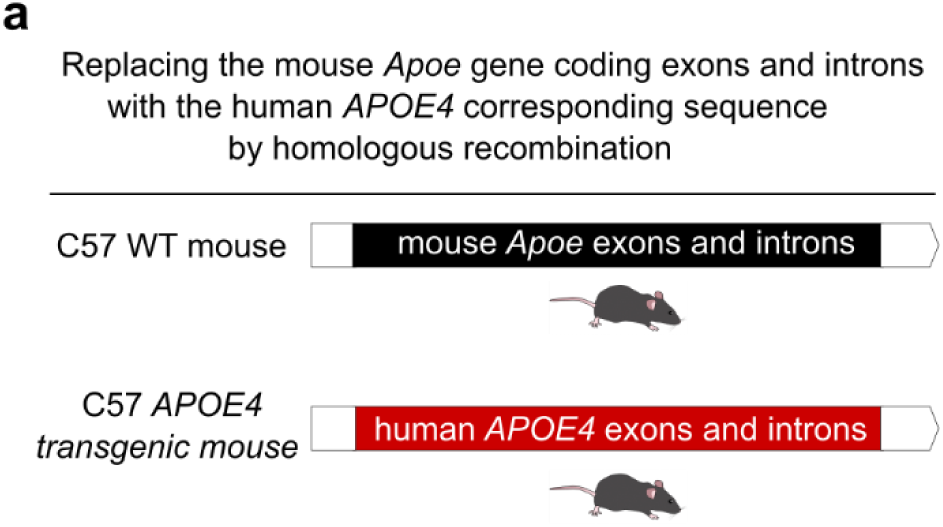
Schematic of the humanized *APOE4* transgenic mice. (a) Schematic showing that the mouse *Apoe* gene (coding exons and introns) was replaced by the human *APOE4* allele by homologous recombination.

**Extended Data Fig. 7.**
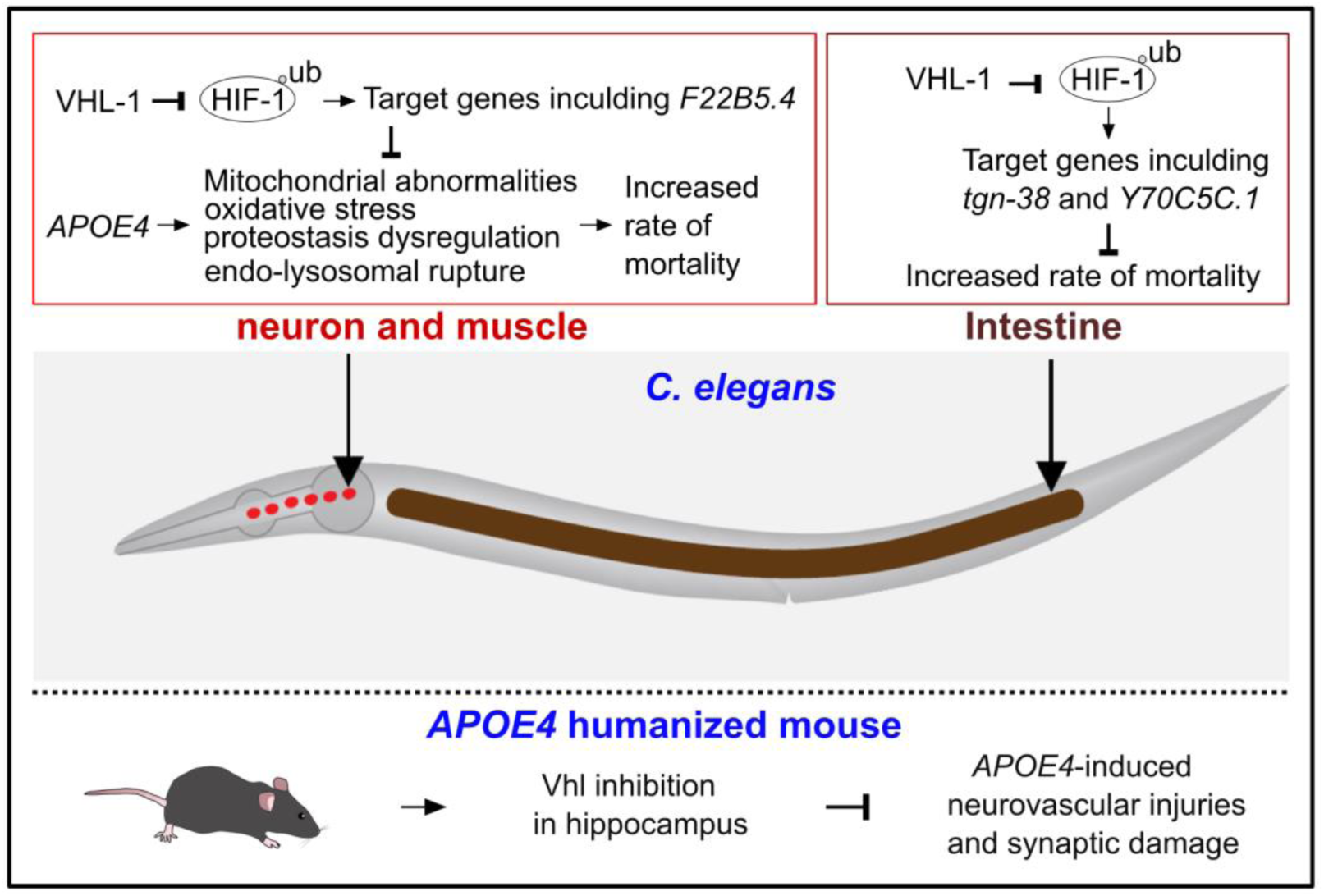
A schematic model.

## Methods

### C. elegans strains

*C. elegans* strains were grown on nematode growth media (NGM) plates seeded with *Escherichia coli* OP50 at 20°C with laboratory standard procedures unless otherwise specified. The N2 Bristol strain was used as the reference wild type. Mutants and integrated transgenes were backcrossed at least 5 times.

Genotypes of strains used are as follows:

Figure 1 and Extended Data Figure 1:

*vhl-1(ok161) X; nIs470 IV; him-5 V,*

*vxIs824 [rab-3p::ND18ApoE4::unc-54 3’UTR + myo-2p::mCherry::unc-54 3’UTR],*

*vhl-1(ok161) X;vxIs824,*

*hpIs376 [unc-25p::tomm20::miniSOG::SL2::RFP],*

*vhl-1(ok161) X; hpIs376,*

*wyIs592[ser-2prom-3p::myr-GFP],*

*vxIs824; wyIs592,*

*vxIs823 [rab-3p::APP::mCherry::unc-54 3’UTR] II. vhl-1(ok161) X; vxIs823,*

*otIs197 [unc-14p::hif-1(P621A) + ttx-3p::RFP],*

*otIs197; vxIs824*

*Ex[rab-3p::APOE3;myo-3p::mcherry] Ex[myo-3p::mcherry]*

Figure 2 and Extended Data Figure 2:

*nIs470 [cysl-2p::GFP + myo-2p::mCherry] IV,*

*hif-1(ia4)*

*vhl-1(ok161);hif-1(ia4);nIs470*

Figure 3 and Extended Data Figure 3:

*dmaIs8 [hsp-16p::GFP; unc-54p::mCherry];him-5(e1490),*

*vxIs824; dmaIs8,*

*rmIs132 [unc-54p::Q35::YFP],*

*vxIs824; rmIs132,*

*rmIs133 [unc-54p::Q40::YFP],*

*vxIs824; rmIs133*

Figure 4 and Extended Data Figure 4:

*vhl-1(ok161) X; vxIs824; wyIs592,*

*hpIs376; wyIs592*

Figure 5 and Extended Data Figure 5:

*dmaIs24[ric-19p::mito::GFP; unc-25p::mCherry],*

*vxIs824; dmaIs24*

*vhl-1(ok161) X; vxIs824; dmaIs24,*

*ceIs56 [unc-129p::ctns-1::mCherry + nlp-21p::Venus + ttx-3p::RFP],*

*vxIs824; ceIs56,*

*ceIs259 [unc-129p::RFP::syn-13 + unc-129p::Venus + ttx-3p::RFP],*

*vxIs824; ceIs259,*

*foxSi41 [dpy-7p::tomm-20::mKate2::HA::tbb-2 3’ UTR] I,*

*vxIs824; foxSi41,*

*xmSi[mai-2::GFP(single copy integration)]*

Figure 6 and Extended Data Figure 6:

*Y70C5C.1(ve718[LoxP + myo-2p::GFP::unc-54 3’ UTR + rps-27p::neoR::unc-54 3’ UTR*

*+ LoxP]) V*.

*tgn-38(gk5592[loxP + myo-2p::GFP::unc-54 3’ UTR + rps-27p::neoR::unc-54 3’ UTR + loxP]) IV*.

*vhl-1(ok161) X; tgn-38(gk5592[loxP + myo-2p::GFP::unc-54 3’ UTR + rps-27p::neoR::unc-54 3’ UTR + loxP]) IV.,*

*dmaEx[ric-19p::F22B5.4; unc54p::mcherry],*

*dmaEx[F22b5.4p::F22B5.4::GFP;unc54p::mcherry],*

*vxIs824; dmaEx[ric-19p::F22B5.4;unc54p::mcherry]*

### Mice and AAV injection

7-month male *APOE4* mice (NM-HU-190002) and C57BL/6 mice (SM-001) were purchased from Shanghai Model Organisms Center, Inc. (Shanghai, China). Animals were maintained on a 12 h light/dark cycle with ad libitum access to food and water. The animal experiments were approved by the Institutional Animal Care and Use Committee of China Pharmaceutical University. For the *in vivo* genetic inhibition of Vhl, mice were randomly divided into the following groups: C57BL/6 + AAV-negative control (NC)-shRNA, C57BL/6 + AAV-*Vhl*-shRNA, *APOE4* + AAV-NC-shRNA, *APOE4* + AAV-*Vhl*-shRNA. AAV (1.0 μl per hippocampus, 1.1 × 10^13^ VG/ml) was injected bilaterally into the hippocampus (from bregma: -2.0 mm AP, ±1.0 mm ML, -2.2 mm DV) of the mice using a stereotaxic apparatus (RWD Life Science Co., Ltd., Shenzhen, China). The cerebral vascular and synaptic assays were performed 30 days after AAV injection.

### Transgenic arrays and strains in *C. elegans*

Transgenic animals that carry non-integrated, extra-chromosomal arrays were generated by co-injecting an injection marker with one to multiple DNA construct at 5– 50 ng/μl. Animals that carry integrated transgenic arrays were generated by UV irradiation (UV Stratalinker2400, Stratagene), followed by outcross at least five times.

### Compound and confocal imaging

Epifluorescence compound microscopes (Leica DM5000 B Automated Upright Microscope System) were used to capture fluorescence images (with a 10× objective lens). Animals of different genotypes and different stages (L1, L2, L4, Day 1 post L4, Day 5 post L4) and different heat treatment were randomly picked and treated with 10 mM sodium azide solution (71290-100MG, Sigma-Aldrich) in M9, aligned on an 2% agarose pad on slides for imaging. The same settings (for bright field: exposure time 1 second, for GFP: exposure time 10-100 seconds) were maintained for the images of all samples. The integrated density (IntDen) of *cysl-2p*::GFP and *hsp16p*::GFP was measured by NIH image program (Fiji image J); averages of mean gray values (three background area of each image randomly selected) were employed for quantification and normalization of *cysl-2p*::GFP and *hsp16p*::GFP. For confocal images, the animals of different genotypes and stages were randomly chosen and treated with 10 mM sodium azide in M9 solution and aligned on an 2% agarose pad on slides and images were acquired using a confocal microscope (Leica TCS SPE) with a 20×, 40× and 63× objective lens, with the same settings maintained for the images of all samples.

### Western blotting

For *C. elegans* samples, stage-synchronized animals for control and experiment groups were picked (n = 50) in 60 μl M9 buffer and lysed directly by adding 20 μl of 4x Laemmli sample buffer (1610747, Bio-Rad) containing 10% of 2-Mercaptoethanol (M6250-100ML, Sigma(v/v)). For cell samples, cultured HEK293T cells were collected by centrifugation followed by adding 60 μl of 1x Laemmli sample buffer containing 10% of 2-Mercaptoethanol. Conditional mediums were collected by centrifugation at 3000*g* for 5 mins at room temperature to remove cell pellets followed by adding 4x Laemmli sample buffer containing 10% of 2-Mercaptoethanol. Protein extracts were denatured at 95 °C for 10 min and separated in 10% SDS-PAGE gels (1610156, Bio-Rad) at 80 V for ∼45 min followed by 110 V for ∼65 min. The proteins were transferred to a nitrocellulose membrane (1620094, Bio-Rad,) at 25 V for 40 mins by Trans-Blot® Turbo™ Transfer System (Bio-Rad). The NC membrane was initially blocked with 5% nonfat milk and 2% BSA (A4503, Sigma (v/v)) in tris buffered saline with 0.1% Tween 20 (93773, Sigma) (TBS-T) at room temperature for 1 h. Proteins of interest were detected using antibodies against pan-actin (4968S, Cell Signaling Technology), pan-APOE (13366S, Cell Signaling Technology) and HIF-1 (14179S, Cell Signaling Technology) in cold room for overnight.

For mouse brain samples, the tissue samples were homogenized, lysed in RIPA buffer (Beyotime, P1003B) containing protease and phosphatase inhibitor (Millipore, 539134 and 524625), and centrifuged at 12,000 rpm for 5 min. Proteins in the supernatant were separated by SDS-PAGE and transferred to PVDF membranes. The membranes were incubated with primary antibodies against Occludin (ABclonal, A2601), Synaptophysin (ABclonal, A19122) or β-actin (ABclonal, AC026) at 4 °C overnight. HRP-conjugated secondary antibodies (Cell Signaling Technology, 7074S) were used, and bands were visualized with ECL chemiluminescence detection kit (Vazyme, E412-01/02) and digitally acquired using Tanon 5200 Multi Chemiluminescent Imaging system (Tanon, Shanghai, China).

### Immunofluorescence

For immunostaining of *C. elegans*, animals were washed with M9 and put in 1.5 ml Eppendorf tubes. The animals were centrifuged for 1 min at 1000*g*, with the liquid removed, and washed again with M9 for three times followed by adding 500 μl ice-cold 4% paraformaldehyde solution for incubation at room temperature for 30 minutes. Fixed animals were washed 3 times by PBS-Tween (0.05%). Animals were centrifuged again with most of the supernatant removed without disturbing the pellet. The pellets were resuspended in 1 mL of 2-mercaptoethanol solution (1ml dH_2_O, 400 μl 0.5 M Tris pH 6.8,15 μl Triton X-100, 76 μl 2-Mercaptoethanol) in the hood followed by incubation at 37°C overnight on a rotator mixer. Samples were then washed 3 times in 1X PBS-Tween (pH 7.2), incubated between each wash with gentle mixing (∼1 hour at room temperature), resuspend in 50 μl of 1X PBS-Tween (pH 7.2) and added with 150 μl of collagenase solution to each tube followed by incubation at 37°C with shaking for 10 min (750 rpm, Eppendorf ThermoMixer F1.5). The samples were then washed 2 times with 1X PBS-Tween (pH 7.2) and 1 time with AbA (40 mL 1X PBS, 200 μl Triton X-100, 0.4 g BSA), resuspended in 200 μl of AbA with primary antibody for anti-pan actin (dilution of 1:1000, 4968S, Cell Signaling Technology) with rocking in cold room for overnight. Samples were washed 3 times with AbA followed by 200 μl of AbA containing Goat anti-Rabbit IgG (H+L) Cross-Adsorbed Secondary Antibody Alexa Fluor® 488 conjugate (dilution of 1:100, A-11008, Thermo Fisher). Samples were kept in the dark and incubated with rocking for overnight at 4°C. Animals were washed 6 times with AbA, incubated with gentle rocking for 1 hr at RT. All animals of different genotypes and conditions were randomly chosen and aligned on a 2% agarose pad on slides and images were acquired using a confocal microscope.

For immunostaining of mouse brain, the brains were fixed in 4% paraformaldehyde solution for 48 h and then embedded in paraffin. Brain sections were cut at 5 μm thickness. Sections were incubated with auto-fluorescence quencher for 5 min to eliminate auto-fluorescence. The sections were incubated with primary antibodies against CD13 (Proteintech, 66211-1-Ig), NeuN (ABclonal, A19086), SMI312 (BioLegend, 837904) for 12 h at 4 °C and subsequently treated with fluorescence-conjugated secondary antibody (Beyotime, A0423 and A0460) for 2 hr at room temperature in the dark. To visualize brain microvessels, sections were incubated with Lycopersicon Esculentum DyLight 488 (Invitrogen, L32470) for 1 hr. The sections were sealed with an anti-fluorescence quencher. Images were acquired using a digital slide scanner (Pannoramic MIDI; 3DHISTECH, Budapest, Hungary). For standard image analysis of fluorescent images, TIFF image files were opened using ImageJ and converted to 8-bit greyscale files. An appropriate threshold value for optimally capturing the intended staining across all conditions in that cohort was determined, and then held constant across all images in the cohort. The areas occupied by CD13+ pericyte on lectin+ brain capillaries, and the areas occupied by the SMI312+ signal were analyzed respectively.

### RNA interference (RNAi)

RNAi were performed by feeding animals with *E. coli* strain HT115 (DE3) expressing double-strand RNA (dsRNA) targeting endogenous genes. Briefly, dsRNA–expressing bacteria were replicated from the Ahringer library to LB plates containing 100 μg/ml ampicillin (BP1760-25, Fisher Scientific) at 37 °C for 16 hrs. Single clone was picked to LB medium containing 100 μg/ml ampicillin at 37 °C for 16 hrs and positive clones (verified by bacteria PCR with pL4440 forward and pL4440 reverse primers) were spread onto NGM plates containing 100 μg/ml ampicillin and 1 mM isopropyl 1-thio-β-Dgalactopyranoside (IPTG, 420322, Millipore) for 24 hrs. Developmentally synchronized embryos from bleaching of gravid adult hermaphrodites were seeded on RNAi plates and grown at 20 °C to L4 followed by transfer to 28 °C for imaging or survival assays.

### qRT-PCR

Animals of synchronized stages (young adults, 24 hrs post L4) were washed off from NGM plates using M9 solution, centrifuged and washed with M9 for three times and subjected to RNA extraction using TissueDisruptor and RNA lysis buffer (Motor unit ‘6’ for 10 seconds and take it out, repeat 3-5 times on ice) and total RNA was extracted following the instructions of the Quick-RNA MiniPrep kit (Zymo Research, R1055) and reverse transcription was performed by SuperScript™ III (18080093, Thermo scientific). Real-time PCR was performed by using ChamQ Universal SYBR qPCR Master Mix (Q711-02, Vazyme) on the Roche LightCycler96 (Roche, 05815916001) system. Ct values of target gene were normalized to measurements of *rps-23* (*C. elegans*) levels. Primers for qRT-PCR were listed in the key resources table.

### RNA-seq

Bleach-synchronized embryos of wild type and *APOE4*-transgenic *C. elegans* (*vxIs824*) were grown to L4 stages with characteristic crescent vulva followed by culture for 24 hours at 25 °C. Animals were washed off from NGM plates using M9, centrifuged and washed with M9 for three times and subjected to RNA extraction using TissueDisruptor and the RNeasy Mini Kit from Qiagen. Three biological replicates were included for WT and *APOE4*. RNA sequencing was performed by BGI American Corporation (DNBseq-G400 platform). An average of 43 million paired reads were generated per sample and the percent of mRNA bases per sample ranged from 46% to 89%. Sequences were aligned to ensemble *C. elegans* genome WBcel235 and read counts per gene were tabulated. All statistical analysis of RNA-seq data was conducted in R v.4.0.5, and count normalization and differential gene expression was performed using the R package DESeq2. Three independent replicates were analyzed for each experiment.

### Thermal resilience and lifespan assays

For thermal resilience assays, animals were cultured under non-starved conditions for at least 2 generations before heat stress assays. (1) For normal NGM thermal resilience assays, synchronized L4 stage animals (n ≥ 50) were picked to new normal NGM plates seeded with OP50 and transferred to 28 °C incubator. (2) For cholesterol free NGM thermal assays, animals were cultured under non-starved conditions for at least one generation on cholesterol free NGM, and synchronized embryos grown up to L4 stage on cholesterol free NGM plates were seeded with OP50 and animals (n ≥ 50) were picked to new cholesterol free NGM plates seeded with OP50 and transferred to 28 °C incubator. (3) For early-life cholesterol free NGM thermal assays, animals were cultured under non-starved conditions for at least one generation on cholesterol free NGM, synchronized embryos grown up to L4 stage on cholesterol free NGM plates seeded with OP50, and animals (n ≥50) were picked to new normal NGM plates seeded with OP50 and transferred to 28 °C incubator. (4) For post L4-life cholesterol free NGM thermal assays, animals were cultured under non-starved conditions on normal NGM, synchronized embryos grown up to L4 stage on normal NGM plates seeded with OP50 and animals (n ≥ 50) were picked to new cholesterol free NGM plates seeded with OP50 and transferred to 28 °C incubator. Animals were scored for survival every 24 hrs. Animals failing to respond to repeated touch of a platinum wire were scored as dead. For lifespan assays, Animals were cultured under non-starved conditions for at least 2 generations before life span assays. (1) For normal NGM life span assay, stage-synchronized L4 stage animals (n ≥ 50) were picked to new NGM plates seeded with OP50 containing 50 μM 5-fluoro-2′-deoxyuridine (FUDR) to prevent embryo growth at 20 °C incubator. (2) For cholesterol free NGM life span assay, animals were cultured under non-starved conditions for at least one generation on cholesterol free NGM, synchronized embryos grown up to L4 stage on cholesterol free NGM plates seeded with OP50 and animals (n ≥ 50) were picked to new cholesterol free NGM plates seeded with OP50 containing 50 μM FUDR and transferred to 20 °C incubator. (3) For early-life cholesterol free NGM life span assay, animals were cultured under non-starved conditions for at least 1 generation on cholesterol free NGM, synchronized embryos grown up to L4 stage on cholesterol free NGM plates seeded with OP50 and animals (n ≥ 50) were picked to new normal NGM plates seeded with OP50 containing 50 μM FUDR and transferred to 20 °C incubator. (4) For early-life cholesterol free and supplementation with NAC diet life span assay, animals were cultured under non-starved conditions for at least 1 generation on cholesterol free NGM, synchronized embryos grown up to L4 stage on cholesterol free and supplementation with 10 mg/ml NAC NGM plates seeded with OP50 and animals (n ≥ 50) were picked to new normal NGM plates seeded with OP50 containing 50 μM FUDR and transferred to 20 °C incubator. Animals were scored for survival per 24 hrs. Animals failing to respond to repeated touch of a platinum wire were scored as dead.

### miniSOG assay

For normal NGM based miniSOG assay, stage-synchronized L4 stage animals (n≥50) were randomly picked to 20 μl M9 solution on the 35 mm dish without lid followed by exposure under 470 nm blue light for 45-90 min under the epifluorescence microscope (SMZ18, Nikon) in the dark room. For early-life cholesterol free NGM based miniSOG assay, animals were cultured under non-starved conditions for at least one generation on cholesterol free NGM, and stage synchronized L4 stage animals (n ≥ 50) were picked to 20 μl M9 solution on the 35 mm dish without lid followed by exposure under 470 nm blue light for 45 min under the epifluorescence microscope (SMZ18, Nikon) in the dark room. For early-life anti-oxidative diet (NAC treatment) based miniSOG assay, animals were cultured under non-starved conditions for at least one generation on normal 60 mm NGM supplemented with 300 ul of 10 mg/ml NAC, stage synchronized L4 stage animals (n ≥ 50) were picked to 20 μl M9 solution on the 35 mm dish without lid followed by exposure under 470 nm blue light for 45 min under the epifluorescence microscope (SMZ18, Nikon) in the dark room. Animals were transferred to NGM plates seeded with OP50. Animals were scored for survival per 1 hr. Animals failing to respond to repeated touch of a platinum wire were scored as dead.

### NAC Compound treatment

For normal NGM thermal resilience, 60 mm dish normal NGM were seeded with of 300 ul N-acetyl cysteine (NAC) at concentration of 1 mg/ml or 10 mg/ml, (1) for early-life with NAC diet, bleached-eggs were transferred to NGM plates supplemented with NAC and grown up to L4 stage at 20 °C incubator followed by transfer to 28 °C incubator, (2) for post-L4 life with NAC diet, L4 stage-synchronized animals from normal NGM plates at 20 °C incubator were picked to NGM plates supplemented with NAC followed by transfer to 28 °C incubator. For early-life cholesterol free and NAC diet thermal resilience, animals were cultured under non-starved conditions for at least one generation on cholesterol free NGM, bleached-embryos were transferred to cholesterol free NGM plates supplement with NAC and grown up to L4 stages at 20 °C incubator followed by pickeing to normal NGM supplemented with 10 mg/ml NAC and transferred to 28 °C incubator. For all NAC diet only-based imaging (*hsp16p::GFP*, *unc54p::Q40::YFP* and PVD::GFP), 60 mm dish normal NGM were seeded with of 300 ul N-acetyl cysteine (NAC) at concentration of 10 mg/ml, bleach-synchronized embryos from normal NGM were transferred to NGM plates supplemented with NAC and grown up to L4 and young adult stages at 20 °C incubator.

For all cholesterol free and NAC diet based imaging (*hsp16p::GFP*, *unc54p::*Q40::YFP), 60 mm dish normal NGM with OP50 were seeded with of 300 ul N-acetyl cysteine (NAC) at concentration of 10 mg/ml, animals were cultured under non-starved conditions for at least one generation on cholesterol free NGM, bleach-synchronized embryos were transferred to cholesterol free NGM plates supplemented with NAC and grown up to L4 and young adult stages at 20 °C incubator. Animals were randomly picked and treated with 10 mM sodium azide in M9 buffer and aligned on a 2% agarose pad on slides for microscopic imaging.

### Animal body size assay

Animals on the normal NGM plates were washed by 1 ml M9 buffer, followed by transfer to 1.5 ml EP tube and addition of 100 μl 5M NaOH, and 200 μl bleach solutions, mixed well and incubated for 4 min (with gentle shaking each mins) at room temperature. The samples were then centrifuged at 850 *g* for 30 seconds to remove the supernatant, followed by 1 ml M9 buffer resuspension, and 850 *g* for 30 seconds for twice. The eggs were resuspended with M9 buffer (200 μl) and transferred to new normal NGM plates or cholesterol free NGM seeded with OP50 and incubated at 20 °C and grown for 48 hrs. Animals were randomly picked and treatment with 10 mM sodium azide in M9 buffer and aligned on a 2% agarose pad on slides for compound microscope imaging. Animals’ relative body size was measured by NIH image program (Fiji image J) based on the area of the worm using freehand lines in the program.

### Genomic DNA damage

For HEK293T cell cultures, parental HEK293T cells and stable-HIF-1 lines were transfected with indicated plasmids for 48 hrs, followed by transfer to 42 °C incubator or 37 °C incubator for 24 hrs. The genomic DNA samples were extracted using Qiagen kits and followed by loading to 1.5% agarose gel for imaging.

### Plasmids

p-mCherry-N1 vector expressing human ApoE2, ApoE3, and ApoE4 were provided by Aparna Lakkaraju (University of California, San Francisco). pLenti PGK Puro vectors expressing stable-HIF-11 (Plasmid #177202, addgene), pMD2.G (Plasmid #12259, addgene) and psPAX2 (Plasmid #12260, addgene) were ordered from Addgene.

### Cell culture and transfection

Human embryonic kidney (HEK) 293T cells (CRL-3216, ATCC) were maintained in Dulbecco’s modified Eagle’s medium with 10% inactive fetal bovine serum (FBS) and penicillin-streptomycin (Gibco, Grand Island, 15140122) at 37 °C supplied with 5% CO_2_ in an incubator (Thermo Fisher Scientific) with a humidified atmosphere. Cells were washed once using PBS and digested with 0.25% trypsin-EDTA (Gibco) at 37 °C for routine passage. HEK 293T cells were transiently transfected with indicated constructs using the lipo2000 (1 mg/ml, LIFE technologies) reagents. The lipo2000/DNA mixture with the ratio of lipo2000 to DNA at 3:1 was incubated for 30 min at room temperature before being added to the HEK 293T cell cultures dropwise.

### Lentivirus and Cell line generation

Lentiviruses were produced by transfecting the HEK293T cells with the pLenti PGK Puro vectors expressing stable-HIF-1 (Plasmid #177202, addgene), and two helper plasmids pMD2.G (Plasmid #12259, addgene) and psPAX2 (Plasmid #12260, addgene), The transfections were carried out using the Polyethylenimine (PEI) method with the ratio at PEI: pLenti PGK Puro: psPAX2: pMD2.G = 18:3:2:1. The lentivirus-containing medium was harvested 72 hrs after transfection and subsequently pre-cleaned with a 3,000 *g* centrifuge for 5 min. The HEK293T cells were incubated with stable-*hif*-1 lentivirus medium with culture medium containing with 5 ug/ml puromycin in a humidified incubator at 37°C with 5% CO_2_. The cultured medium was changed after 24 hrs with fresh medium containing 5 ug/ml puromycin. The stable-HIF-1 positive HEK293T cell lines were maintained with medium containing with 5 ug/ml puromycin.

### Mammalian cell thermal resilience assay

For thermal resilience assay, mock control and transfected HEK293T cells or stable-hif-1 HEK293T cell line (48 h) in 24 well plates were placed in a culture incubator with an ambient temperature at 42°C and humidified 5% CO_2_ for 8-24 hrs followed by cell death assay, genomic DNA extract or imaging with 4% PFA treatment for 12 min at room temperature. For cell death assay, the collected cells were resuspended with 100 μl buffer with addition of 0.1 μl Sytox blue (Thermo Fisher Scientific) for an additional 15 min at room temperature. 25 ul of incubated cells were loaded into ArthurTM cell analysis slide (Nanoentek, AC0050). The fluorescence intensity was measured for individual cells using automated cytometry (ArthureTM image-based cytometer, Nanoentek, AT1000) as viability assay. The 190 RFU (Fluorescence) threshold and cell size min 5 to max 25 were used for cell death analysis and quantification.

### Conditional medium and cell lysis treatment with *C. elegans*

For conditional medium treatment with *C. elegans*, the conditional medium of HEK293T cells and stable-hif-1 HEK293T cell line were collected to 1.5 ml EP tubes, and subsequently pre-cleaned with a 3,000 *g* centrifuge for 5 mins to remove cell pellets. 60 mm dish normal NGM were seeded with of 300 ul defined conditional medium. 24 hrs later, bleach-synchronized embryos were transferred to NGM containing with conditional medium and grown up to L4 stage at 20 °C incubator followed by transfer to 28 °C incubator. For HEK293T cell lysate treatment, HEK293T cells and stable-hif-1 cells transfected with mCherry or APOE4::mCherry were collected followed by lysis using lysis buffer (50 mM Hepes, pH 7.4, 100 mM NaCl, 50 mM KCl, 2 mM CaCl_2_, 2 mM MgCl_2_, 1 mM PMSF, one tablet of protease inhibitor cocktail per 50 ml buffer containing 0.5 % Triton X-100.) Insoluble material was removed by centrifugation (10,000 *g* for 10 min), and the supernatant was used for seeding onto 60 mm dish normal NGM, bleach-synchronized embryos were transferred to NGM containing the conditional medium and grown up to L4 stage at 20 °C incubator followed by transfer to 28 °C incubator.

### Evans blue leakage

Evans blue dye (2% in saline, 4 ml/kg body weight, Macklin, E6135-1g) was injected intravenously into mice. Mice were transcardially perfused with saline, and brain tissues were harvested after 1 h. The brain tissues were subsequently homogenized in 2 ml of 50% trichloroacetic acid, and centrifuged at 12000 rpm for 30 min. The supernatants were measured for optical density at 620 nm using a Spectramax 190 Microplate Reader (Molecular Devices, San Jose, USA).

### Statistical analysis

Data were analyzed using GraphPad Prism 9.2.0 Software (Graphpad, San Diego, CA) and presented as means ± S.D. unless otherwise specified, with P values calculated by unpaired two-tailed t-tests (comparisons between two groups), one-way ANOVA (comparisons across more than two groups) and two-way ANOVA (interaction between genotype and treatment), with post-hoc Tukey HSD and Bonferroni’s corrections. The lifespan assay was quantified using Kaplan–Meier lifespan analysis, and P values were calculated using the log-rank test.

## Acknowledgment

Some strains were provided by the *Caenorhabditis* Genetics Center (CGC), which is funded by the NIH Office of Research Infrastructure Programs (P40 OD010440), by Drs. Kang Shen, Rosa Navarro, and Jo Anne Powell-Coffman. The work was supported by NIH grants (R35GM139618 to D.K.M., RF1AG057355 and R21OD032463 to J.T.P.), BARI Investigator Award (D.K.M.), UCSF PBBR New Frontier Research Award (D.K.M.), National Natural Science Foundation of China (No. 82173728, S.C.), UCSF CIRM Scholars Training Program EDUC4-12812 (W.J.), Schmidt Science Fellowship (B.Y.W.) and Cancer Research Irvington postdoctoral fellowship (B.Y.W.).

## Author contributions

W.J. and D.K.M. designed, performed and analyzed the *C. elegans* and cell culture experiments, contributed to project conceptualization and wrote the manuscript. Y.C., Y.X., Y.J. and S.C. designed, performed and analyzed the mouse experiments, contributed to project conceptualization and wrote the manuscript. B.Y.W. analyzed the RNA-seq data. N.S., M.Z., and J.P. contributed to research materials, project conceptualization and editing manuscript.

